# Visual imagery of familiar people and places in category selective cortex

**DOI:** 10.1101/2024.09.13.612846

**Authors:** Catriona L. Scrivener, Jessica A. Teed, Edward H. Silson

**Affiliations:** School of Philosophy, Psychology and Language Sciences, University of Edinburgh, Edinburgh, UK; School of Psychology and Neuroscience, University of Glasgow, Glasgow, UK

**Keywords:** mental imagery, EEG-MRI fusion, category selectivity

## Abstract

Visual imagery is a dynamic process recruiting a network of brain regions. We used electroencephalography (EEG) and functional magnetic resonance imaging (fMRI) fusion to investigate the dynamics of category selective imagery in medial parietal cortex (MPC), ventral temporal cortex (VTC), and primary visual cortex (V1). Subjects attended separate EEG and fMRI sessions where they created mental images of personally familiar people and place stimuli. The fMRI contrast comparing people and place imagery replicated previous findings of category-selectivity in the medial parietal cortex. In addition, greater activity for places was found in the ventral and lateral place memory areas, the frontal eye fields, the inferior temporal sulcus, and the intraparietal sulcus. In contrast, greater activity for people was found in the fusiform face area and the right posterior superior temporal sulcus. Using multivariate decoding analysis in fMRI, we could decode individual stimuli within the preferred category in VTC. A more complex pattern emerged in MPC, which represented information that was not restricted to the preferred category. We were also able to decode category and individual stimuli in the EEG data. EEG-fMRI fusion indicated similar timings in MPC and VTC activity during imagery. However, in the VTC, fusion was higher in place selective regions during an early time window, and higher in face selective regions in a later time window. In contrast, fusion correlations in V1 occurred later during the imagery period, possibly reflecting the top-down progression of mental imagery from category-selective regions to primary visual cortex.

## 1. Introduction

The ability to picture something in your ‘minds-eye’ in the absence of an external stimulus is referred to as mental imagery; an everyday task with surprising variation across individuals (Zeman, 2024). One theory of visual mental imagery describes the phenomenon as perception in reverse (Pearson, 2019). Perception of visual stimuli activates a hierarchy of brain regions, extending from early visual cortex into dorsal and ventral streams, further recruiting parietal and frontal regions in a task dependent manner. Visual mental imagery might then be characterised by activity in similar regions but in the reverse order, culminating in a reinstatement of ‘perception-like’ activity in primary visual cortex (V1) (Dijkstra et al., 2017). This hypothesis has led to a focus of exploration in V1, with groups searching for similar functional magnetic resonance imaging (fMRI) blood oxygenation level dependent (BOLD) activity or representational patterns across perception and imagery of the same stimuli. There has been some success in this approach for visual imagery of simple stimuli such as gratings, letters, and objects (Albers et al., 2013; Lee et al., 2012; Ragni et al., 2020). However, successful comparison of perception and imagery in V1 may be influenced by the stimuli used and the task given to participants (Dijkstra et al., 2019; Kosslyn & Thompson, 2003).

An alternative theory is that mental imagery recruits a flexible network of category-selective, memory, and semantic processing regions that vary with the specific imagery content and task performed (Spagna et al., 2024). It is plausible that regions of the brain outside of early visual cortex are also involved in generating mental images, especially for images that require other cognitive processes such as engaging memory representations or mental rotation (Dijkstra et al., 2019; Zeman, 2024). One example of this is the visual imagery of specific people and places, which has been found to activate several regions within medial parietal cortex (MPC) (Silson, Steel, et al., 2019). These regions display category-selectivity, with increased BOLD activation during imagery of stimuli from their preferred category (people or places) that is elevated when the stimuli are personally familiar. Several regions in the ventral temporal cortex (VTC) are also active during imagery, including the parahippocampal place area (PPA, Epstein & Kanwisher, 1998) and fusiform face area (FFA, Kanwisher et al., 1997), despite mostly being studied in the context of visual perception. The link between these category-selective regions in medial parietal and ventral temporal cortex is indicated by their increased resting-state connectivity. For example, the anterior parahippocampal place area (aPPA, Baldassano et al., 2016) is most functionally connected with the largest place selective region in medial parietal cortex (Places 1; also referred to as ventral medial parietal cortex (MPCv) in Silson et al., 2019), whereas the anterior fusiform face area (FFA2) is most functionally connected with the analogous face selective region (People 1; also referred to as dorsal medial parietal cortex (MPCd) in Silson et al., 2019).

Although these category-selective regions in medial parietal and ventral temporal cortex are all engaged during imagery of people and places, it is currently unclear if they contribute similarly to the generation of visual imagery or if they play distinct roles. During perception, medial parietal regions typically demonstrate a decrease in activation, which is related to their proximity to the default mode network (Raichle et al., 2001; Silson, Steel, et al., 2019) and differentiates them from their ventral temporal counterparts. It has also been suggested that category-selective regions within medial parietal cortex form part of a semantic network, based on their overlap with regions representing the semantic content of visual images (Koslov et al., 2024). Other work highlights the role of the posterior parietal cortex in supporting episodic memory (Bainbridge & Baker, 2022; Ranganath & Ritchey, 2012; Ritchey & Cooper, 2020). It is therefore plausible that the activity of medial parietal and ventral temporal regions during imagery will differ, although it is currently unclear exactly how. Given that these ROIs are distributed along a ventral-dorsal axis within cortex, it is possible that their activity may emerge at different timepoints during mental imagery, and that this may differ from V1.

Prior work using electroencephalography (EEG) and magnetoencephalography (MEG) provides evidence for the time course of visual mental imagery across the brain as a whole. Significant decoding of imagery stimuli has been found to emerge between 540ms (Dijkstra et al., 2017) and 1000ms (Corriveau et al., 2022) post imagery onset, with sustained representations throughout the imagery period. Further, cross-decoding using perceptual responses to the same stimuli suggests that perception data at 150ms, 200ms, and 300ms is most predictive of imagery representations (Dijkstra et al., 2017; Xie et al., 2020), and that ‘perception-like’ imagery emerges from 400ms (Dijkstra et al., 2017) or 700ms (Xie et al., 2020) onwards (the differences in the exact timings across these studies are likely caused by variation in design and analysis choices). While these results provide valuable information regarding the emergence of visual imagery through time, it is difficult to assess the source of this information in the brain, or how different regions may contribute to these imagery representations. To address this, we used an EEG-fMRI fusion approach (Cichy & Oliva, 2020) to examine how the time course of visual mental imagery recorded in EEG relates to activity across several key regions; medial parietal cortex (People and Place regions), ventral temporal cortex (PPA and FFA), and V1. We hypothesised that we may see different timings across medial parietal and ventral temporal category-selective ROIs, reflecting their different roles in the generation of mental images. We also hypothesised that we would find evidence of a top-down progression of mental imagery, with representations of people and places emerging earlier in higher-level visual cortex than in V1.

## 2. Methods

### 2.1 Experimental procedure

A total of 12 subjects participated in two experimental sessions, one with EEG recording and the other with fMRI (mean age = 27, 3 males). Ethical approval was provided by the University of Edinburgh (420-2122/5 and 420-2122/6). All participants gave informed consent and were given monetary compensation for their time. Separate EEG and MRI recordings were appropriate for this experiment as we did not intend to look at trial-by-trial fluctuations in a time-dependent manner (Scrivener, 2021). Instead, we planned to capture the shared information between conditions using a representational similarity analysis (RSA) approach (Nili et al., 2014). Before attending their first session, subjects were asked to provide the researchers with images of six personally familiar people and six personally familiar places that they were confident they could easily create a mental image of. We asked subjects to save these images with a name that would serve as a cue for them to imagine the stimuli, such as ‘Sophie’, or ‘Office’. For the exact details of our request for stimuli, refer to the supplementary material (S1).

During the EEG session, subjects completed two runs of a perception task lasting 4.5 minutes each, where each of the 12 stimuli were presented 25 times per run in a randomised order (50 total repetitions of each stimuli, stimuli presented for 500ms, with a jittered inter-trial interval between 300 and 500ms). Stimuli were presented at 12 by 12 degrees of visual angle. Subjects were asked to look at a central fixation cross and press ‘space’ when it turned from green to red during a random subset of trials. They then completed four runs of the imagery task lasting 6.8 minutes each **(Figure 1)**, where each imagery cue was presented 10 times per run in a randomised order (40 total repetitions per stimuli). One subject only completed 3 out of the 4 imagery runs due to a technical error during the last run. Subjects were specifically instructed to create a mental image of the picture they had seen during the perception runs. This was an attempt to standardise their mental imagery across repetitions of the same stimuli, and to make it clear that they should be thinking of the images rather than about general aspects of the people and places (as much as possible). Subjects were also instructed to keep their eyes open and remain fixated on the cross. In this manuscript we focus on analysis of the imagery data only, leaving consideration of the perception data for future work.

**Figure 1.**
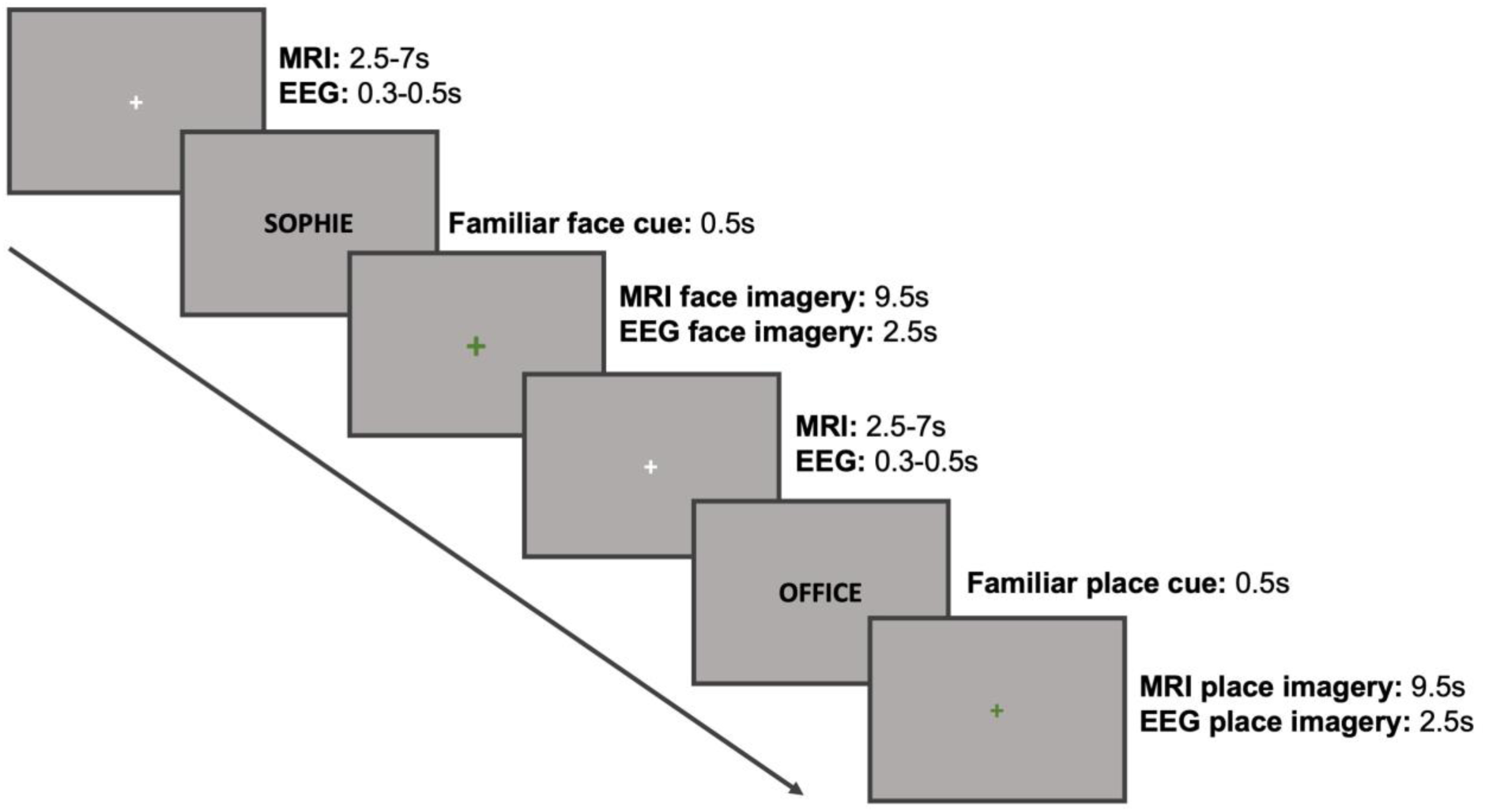
Mental imagery paradigm during EEG and fMRI sessions. Each trial began with a white fixation cross with a jittered duration. Subjects were then presented with a visual cue for 0.5 seconds telling them which of their images they should imagine. This was followed by a green fixation cross which indicated that they should visualise the cued image.

During the MRI session, subjects completed two hours of scanning with a break in between (ranging from 15-60 minutes, depending on scanner timetabling constraints). In session 1, we acquired T1 weighted and T2 weighted structural scans. These were followed by three runs of the perception task, lasting 4.4 minutes each, where each of the 12 stimuli were presented four times per run (12 total repetitions, stimuli presented for 500ms, with a jittered inter-trial interval between 2500 to 7000ms). Note that we do not use the perception runs at all in the current manuscript. This was followed by four functional runs of population receptive field (pRF) mapping (Dumoulin & Wandell, 2008), lasting six minutes each. For pRF mapping we used a bar aperture that made eight sweeps through the visual field (18 positions, 36s per sweep). Scene fragments from five images were presented inside the bar aperture during each position (400ms per scene), sampled randomly from a list of 90 possible scenes (Silson et al., 2015). During both pRF and perception tasks, subjects were asked to fixate a central cross and respond with a right-hand button press when it turned from green to red during a random subset of trials.

In session 2, we acquired a second T1w structural scan to facilitate across session alignment and segmentation if needed. This was followed by two runs of a scene/face localiser task, lasting 5 minutes each. Subjects viewed 18 blocks of 20 scene or face stimuli that lasted 16s each (300ms on, 500ms off), and were asked to complete a one-back task by making a right-hand button press if the same image was repeated sequentially (twice per block). Finally, they completed six runs of the imagery task, lasting six minutes each. Each imagery cue was presented twice per run (12 total repetitions). One participant only completed four of the six imagery runs. See **Figure 1** for the stimulus timing information during imagery.

At the end of both EEG and MRI sessions, subjects were asked to complete the same short questionnaire regarding each stimulus presented during the session. This included an estimate of the overall vividness of their imagery per stimulus (1 = no imagery, 5 = moderately strong imagery, 10 = very clear imagery). All subjects also completed a paper version of the Vividness of Visual Imagery Questionnaire (VVIQ) during their first experimental session.

### 2.2 Visual properties of the stimuli

We ran the images provided to us by the subjects through two image computable models to provide quantification of the visual differences between them. First, we used the gist model, which characterises each image by their correspondence with Gabor filters of varying spatial frequency and orientation (Oliva & Torralba, 2006). We also used the LGN model, which describes the spatial coherence (SC) and contrast energy (CE) within the image (Groen et al., 2013). Of course, these quantifications will only partially relate to the imagery of these stimuli, which may not be a faithful replication of the original image. However, we hoped that these statistics might capture some of the underlying variation in activation across stimuli.

### 2.3 EEG acquisition and pre-processing

EEG data was recorded in ActiView using a BioSemi ActiveTwo AD-box amplifier and 64 pin-type active electrodes. Two additional flat-type free electrodes were secured to the left and right mastoids underneath the EEG cap. Data was acquired at a sampling rate of 512Hz. A technical error meant that the data for one subject was recorded at 128Hz. Their data was up-sampled in MATLAB for plotting purposes only (interp). Subjects were seated approximately 57cm from a computer monitor that displayed the task (size 1920 x 1080). EEG data was pre-processed using EEGLab (2022.0, Delorme & Makeig, 2004) in MATLAB (2023b). The data was re-referenced to the left and right mastoid electrodes (pop_biosig) and then filtered using a Hamming window bandpass filter between 0.1 and 40Hz (pop_eegfiltnew). We then downsampled the data recorded at 512Hz to 256Hz to reduce computation time (pop_resample). Noisy channels were interpolated using a spherical interpolation (pop_interp) for two subjects only (005, 3 electrodes; 006, 7 electrodes). Noisy trials were identified by eye before removal, using a conservative criterion of any time period with large movement or muscle artifacts that affected more than 50% of the channels. On average, 12 of the recorded trials were removed per subject across the whole EEG session (M = 12, SD = 12). For the imagery data, the minimum number of trials remaining for any given stimulus was 20, and on average there were 38 trials included per stimulus (calculated across all subjects and stimuli). Break periods were also removed as they often contained large artifacts. ICA was then performed to identify blink and lateralized eye movements (pop_runica, runica with extended informax). One blink component was removed for all subjects, and one lateralised eye movement was removed for 11 of the 12 subjects (pop_subcomp) after manual inspection of the component topologies and timeseries. Epochs were created from −200ms:3000ms in the imagery runs in relation to the onset of the cue (pop_epoch). Baseline correction was performed using the timepoints −200ms:0ms (pop_rmbase).

### 2.4 MRI acquisition

MRI scans were acquired using a Siemens 3T Prisma scanner and a 32-channel head coil at the University of Edinburgh Imaging Facility RIE, Edinburgh Royal Infirmary. We acquired two types of structural image; T1 weighted (TR = 2.5s, TE = 4.37ms, flip angle = 7 deg, FOV = 256mm x 256mm x 192mm, resolution = 1mm isotropic, acceleration factor = 3), and T2 weighted (TR = 3.2s, TE = 408ms, flip angle = 9 deg, FOV = 256mm x 240mm x 192mm, resolution = 0.9mm isotropic, acceleration factor = 2). Functional scans were acquired using a multiecho multiband echo planar imaging sequence (TR = 2, TEs = 14.6ms, 35.79ms, 56.98ms, MB factor = 2, acceleration factor = 2, 48 interleaved slices, phase encoding anterior to posterior, transversal orientation, slice thickness = 2.7mm, voxel size = 2.7mm x 2.7mm, distance factor = 0%, flip angle = 70 degrees).

### 2.5 MRI pre-processing

MRI scans were processed using AFNI, SPM, Freesurfer, and SUMA (R. W. Cox, 1996; Saad & Reynolds, 2012). Before pre-processing, the origin of all MRI data was manually reset to the anterior commissure (AC) using SPM display (SPM12). For the structural images, the AC origin was defined in the T1w image and the transformation applied to the T2w image, to ensure that they were aligned to each other. For the EPIs in each run, the origin was defined in the first TR of the second echo of the first run, and applied to all other echoes and scans in the same session. This procedure was to facilitate co-registration of EPIs and structural scans across the two sessions, given that the origins of the acquired data could be far apart in space.

During pre-processing, dummy scans were first removed from the start of each run (AFNI 3dTcat). Large deviations in signal were removed (3dDespike) followed by slice time correction (3dTshift), aligning each slice with a time offset of zero. The skull was removed from the first echo 1 scan (TE = 14.6ms) and used to create a brain mask (3dSkullStrip and 3dAutomask), as this echo contained the most signal. The first echo 2 scan (TE = 35.79ms) was used as a base for motion correction and registration with the T1 structural scan (3dbucket), as this echo was the most similar to standard EPI acquisition. For session 1 we used the pRF scans as a base and for session 2 we used the scene/face localiser scans. Motion parameters were estimated for the echo 2 scans (3dVolreg) and applied to the other echos (3dAllineate). After completing the standard pre-processing, the data were processed using Tedana (DuPre et al., 2021; Kundu et al., 2012, 2013) to denoise the multi-echo scans (version 0.0.12, using default options). The Tedana optimally combined and denoised output was then scaled. To do this, we divided the signal in each voxel by its mean value and multiplied the signal by 100 (3dTstat and 3dcalc). This means that the fMRI values can be interpreted as a percentage of the mean signal, and effect estimates can be viewed as percentage change from baseline (Chen et al., 2017). For the pRF scans, an average was then calculated across runs to leave a single time series for further analysis.

The session 1 structural scans were aligned to the functional data collected in session 1 (align_epi_anat) with a multi-cost function (including LPC, MNI, and LPA) and manually checked for accuracy. For all subjects we used the default LPC method output (Localised Pearson Correlation), as this worked sufficiently well. Functional data collected in session 2 was aligned to the session 1 structural to ensure that all functional data had the same alignment. Freesurfer reconstructions were estimated using both the T1 and T2 scans (recon-all) from session 1, and the output used to create surfaces readable in SUMA (SUMA_Make_Spec_FS). For one subject we also included the session 2 structural to aid the reconstruction, as there were movement artifacts in the session 1 structural. The SUMA structural was then aligned to the session 1 experimental structural to ensure alignment with the functional images (SUMA_AlignToExperiment). Surface based analyses were conducted using the SUMA standard cortical surface (std.141).

### 2.6 EEG analysis

#### 2.6.1 Multivariate timeseries decoding

Decoding analyses were performed using CoSMoMVPA (Oosterhof et al., 2016) and custom MATLAB scripts. A linear support vector machine (libSVM) classifier was trained to distinguish between all individual stimuli (1-12) across all channels and timepoints. For category-level decoding, we took an average of all across-category decoding timeseries (e.g. person1 versus place1 etc.). For stimulus-level decoding, we took an average of all within-category decoding timeseries (e.g. person1 versus person2 etc.). We used a leave-one-pseudotrial-out cross-validation approach, where pseudotrials were created using chunks of trials that were averaged together before classification. We chose to use averages of 5 trials. This decision was based on the total number of trials available for each condition, and on previous research evaluating the trade-off between reduced signal-to-noise provided by averaging with the subsequent increases in between-subject variation (Grootswagers et al., 2016; Scrivener et al., 2023). To ensure that the results were not dependent on a specific division of trials, we repeated the random allocation of single trials to pseudotrial chunks 50 times per subject and took the average across all iterations. The choice of 50 iterations was to balance sensitivity with computation time. For each subject, we used the average across these 50 iterations. Classification was run using a moving time-window of 3 samples to increase the number of datapoints available to the classifier. To assess the stability of category representation across time (person or place), we also ran a time-generalised analysis (King & Dehaene, 2014). Here the classifier was trained and tested across all pairs of time points. For this method implemented in CoSMoMVPA, the trials are randomly divided into 2 chunks. Given the increased computation required for time-generalised analysis, we only ran 10 iterations of the chunking procedure. Time points with decoding significantly above chance were assessed using Bayesian t-test calculated using the ‘bayesFactor Toolbox’ in MATLAB (https://github.com/klabhub/bayesFactor). This was implemented using a half-Cauchy prior with a default width of 0.707 and a range of 0.5 to increase the detection of small effects under the null hypothesis (Kass & Raftery, 1995; Morey & Wagenmakers, 2014; Rouder et al., 2012; Teichmann et al., 2022).

### 2.7 MRI analysis

#### 2.7.1 Defining regions of interest

In this work, we used a combination of subject-specific ROIs defined using functional scene/face localiser scans (FFA1/2, a/pPPA), subject-specific ROIs defined using population receptive field mapping scans (V1), and a-priori defined ROIs taken from previous work (people and place memory areas in the medial parietal cortex; Silson et al., 2019). We chose to use subject-specific ROIs where possible given their increased sensitivity to detect group level effects (Nieto-Castañón & Fedorenko, 2012). However, we opted to use previously defined ROIs for the medial parietal cortex, as we did not have separate localiser scans for these ROIs and wanted to avoid issues of circularity in our analysis (Kriegeskorte et al., 2009).

To localise the PPA and FFA, a general linear model was estimated for the scene/face localiser scans using a block design with a 16 second GAM basis function (GAM: 8.6, .547, 16, 3dDeconvolve and 3dREMLfit). The output of the model was then projected onto the SUMA cortical surface (3dVol2Surf), and smoothed with a FWHM of 2mm (SurfSmooth). The PPA and FFA ROIs were drawn manually for each subject on the surface (SUMA draw ROI) after thresholding the contrast of scenes versus faces at t > 3.5 (height-defining threshold, p<0.0001, uncorrected). PPA and FFA ROIs were divided into anterior and posterior portions, either by identifying two separate peaks in the t-maps, or by dividing a single large cluster equally along the posterior-anterior axis (Baldassano et al., 2016; Silson, Steel, et al., 2019). Not all ROIs were identifiable across all subjects (left hemisphere: FFA1 = 10, FFA2 = 12, pPPA = 9, aPPA = 12; right hemisphere: FFA1 = 10, FFA2 = 11, pPPA = 12, aPPA = 12). Our surface-based ROIs were also converted to volume masks by projecting them from the surface to the volume, using the same parameters that were used to project the scene/face localiser data to the surface (3dSurf2Vol). A downside to this approach is that the ROIs varied in size across subjects. This may influence the decoding outcomes, given that more data generally facilitates higher decoding accuracies (D. D. Cox & Savoy, 2003; Gardumi et al., 2016). However, all of the subject-specific ROIs were sufficiently large, with more than ∼200 nodes (see supplementary material S2), and with no systematic differences across subjects in ROI size (i.e., the smallest ROIs were not all in same subject).

Medial parietal ROIs, also defined on the surface, were taken from previous work (Silson et al., 2019). Here, two ROIs on the medial parietal surface with increased BOLD activation during imagery of place stimuli were named as Places1 (left hemisphere = 2587 nodes, right hemisphere = 2766) and Places2 (LH = 651 nodes, RH = 1050). A further two ROIs with higher activation for people stimuli were named People1 (LH = 1319 nodes, RH = 1455) and People2 (LH = 136 nodes, RH = 215). We compared the locations of our group level results in the medial parietal cortex with the ROIs taken from Silson et al. (2019) by calculating the dice-coefficient for each ROI (2*Overlap between ROIs / size of ROI A + size ROI B). The dice-coefficient has a range between 0 and 1, with 1 indicating a perfect overlap.

Primary visual cortex (V1) was defined for each subject individually using population receptive field mapping. Population receptive fields were estimated using AFNI’s non-linear fitting algorithm (3dNLfim) and the GAM basis function (Silson et al., 2015). The outputs were used to delineate subject-specific retinotopic maps which were drawn manually on the SUMA standard (std.141) surface using the polar angle and eccentricity estimates. V1 was further divided into foveal and peripheral portions using the eccentricity output from the pRF modelling. We defined foveal as all V1 nodes with eccentricity values up to 4 degrees of visual angle, and peripheral as the remaining nodes within the V1 ROI (Wandell et al., 2007). ROIs were converted into 1D files (ROI2dataset) to facilitate future node selection.

#### 2.7.2 Mental imagery univariate activation

The activity associated with each stimulus in the imagery scans was deconvolved using a BLOCK basis function (BLOCK(9.5,1), 3dDeconvolve and 3dREMLfit) aligned to the onset of the imagery period. Estimates of subject motion were included as regressors of no interest. We ran separate stimuli and category level deconvolutions, producing either one beta value per category (people or place), or one per stimulus (stimuli 1 to 12). In version one of the stimulus level deconvolutions, the six runs were concatenated and each stimulus regressor included 12 onsets (two from each run). The output of the model was projected onto the SUMA standard cortical surface (3dVol2Surf), and the data within each subject specific ROI was extracted for further analysis (pPPA, aPPA, FFA1, FFA2, Places1, People1, Places2, People2). In a second version of the deconvolutions, the runs were modelled separately so that we could check for an effect of run, and to facilitate a leave-one-run-out cross-validation approach for decoding. Group level analysis was conducted on the surface data using a linear mixed model (3dLME). We included subject as a random effect and modelled the main effect of category (people vs. places), the main effect of run (runs 1 to 6), and an interaction between category and run effects. Activation was thresholded at p < .74^-4^ (q < .0094) for the main effect of category, corresponding to a minimum F value of 12. This was informed by prior work (Silson, Steel, et al., 2019). Note that we did not use any smoothing or any cluster-based corrections.

#### 2.7.3 ROI univariate analysis

We entered the averaged beta values per ROI from the category deconvolution into a linear mixed model with factors Region and Category (people and places), keeping the medial parietal and ventral temporal regions separate and averaging over left and right hemisphere (R studio package v1.3, lme4 v27.1, lmerTest v3.1). Subject was entered as a random factor. Univariate analysis including Hemisphere as a factor can be found in the supplementary material (S5). As the timeseries were scaled, the beta values can be interpreted as BOLD percent signal change relative to the voxel baseline (Chen et al., 2017). We also extracted the average contrast beta values for each ROI (places versus people). We used both Frequentist and Bayesian t-tests to identify category selectivity across our ROIs; i.e. contrast values significantly different from zero. Frequentist p-values were corrected for multiple comparisons using the Benjamin-Hochberg procedure (Benjamini & Hochberg, 1995; Yekutieli & Benjamini, 1999) to control the false discovery rate (FDR correction). We report the original p-values as well as the FDR adjusted thresholds used to determine the significance. See the supplementary material for the average timeseries correlations across ROIs (S3).

#### 2.7.4 ROI decoding analysis

ROI decoding using the beta values was implemented in CoSMoMVPA (Oosterhof et al., 2016) using a leave-one-run-out cross-validation approach and a support vector machine classifier (libSVM). To mimic the ROI univariate analysis, we first decoded the stimulus category across our ROIs (person or place). Next, we decoded the stimulus identity separately within each category (person 1 or person 2 etc.). For this we took the average decoding across all pairs of stimuli, meaning that chance level decoding was 50%. Decoding analyses were run in the volume and in each hemisphere separately. We entered the average decoding value across hemispheres for each ROI and each subject into a linear mixed model with factors Region and Category, keeping the medial parietal and ventral temporal regions separate (R studio package v1.3, lme4 v27.1, lmerTest v3.1). Subject was entered as a random factor. Multivariate analysis including Hemisphere as a factor can be found in the supplementary material (S8). Post-hoc tests for decoding above chance within each ROI were assessed using both Frequentist and Bayesian one-way t-tests. Frequentist p-values were corrected for multiple comparisons using the Benjamin-Hochberg procedure (Benjamini & Hochberg, 1995; Yekutieli & Benjamini, 1999) to control the false discovery rate (FDR correction). We report the original p-values as well as the FDR adjusted thresholds used to determine the significance.

#### 2.7.5 EEG-fMRI fusion

We implemented EEG-fMRI fusion using a representational dissimilarity analysis framework (Cichy & Oliva, 2020). The aim of this EEG-fMRI fusion approach is to identify when in time the representational structure across a set of stimuli in a given fMRI region of interest is matched in the EEG time course. In other words, EEG-fMRI fusion timeseries indicate *when* in time the same pattern of responses across conditions is seen in both EEG and fMRI signals. The focus of this approach is therefore to identify the shared information across EEG and fMRI data sets condition-by-condition, rather than trial-by-trial. One benefit of this RSA fusion approach is that the signal is extracted away from the primary EEG and fMRI data formats, which vary considerably in their respective spatial and temporal resolutions (Scrivener, 2021). It is worth noting a downside to this approach; if two fMRI regions represent a stimulus set in a very similar way then they will have correlated similarity matrices and therefore similar fusion timeseries, even if this representation emerged at different points in time. However, it is generally accepted that information travelling across regions is transformed non-linearly, meaning that the representational format will likely differ across the brain, leading to differences in the fusion timeseries (Cichy & Oliva, 2020).

In the fMRI data, we first constructed separate representational dissimilarity matrices (RDMs) for each ROI, in each hemisphere and for each subject. These were constructed using the imagery deconvolution where each stimulus was modelled with one regressor across all runs (see section 2.7.2). The beta values estimated for each of the 12 stimuli across the ROI nodes extracted from the surface data were used to construct 12×12 RDMs (1-Pearson’s correlation r values) ordered by their category (people then places). This resulted in 18 fMRI RDMs per subject (9 ROIs, pPPA, aPPA, FFA1, FFA2, Places1, Places2, People1, People2, & V1, across 2 hemispheres).

Similar RDMs were also constructed for the EEG imagery data. First, an average was calculated across all trials for each of the 12 stimuli. At each time point, RDMs were then constructed using the activity across all sensors for each stimulus (1-Pearson’s correlation r values). These 12×12 RDMs had the same order and structure as the corresponding fMRI RDMs for each subject (people then places).

To calculate the similarity between the stimulus representation across time and space, we then correlated (Spearman’s rho) the EEG RDMs at each time point with the corresponding fMRI RDM for each ROI in each hemisphere. We then averaged the fusion correlations across hemispheres for each subject. The comparison of fusion correlations across hemispheres can be found in the supplementary material (S9). Time points with fusion correlations above zero were assessed using Bayesian t-tests as in the EEG decoding analyses.

To directly compare the fusion correlation timings across ROIs and conditions, we also calculated a series of difference waves. First, to compare the EEG-fMRI fusion in category selective ROIs with V1, we subtracted the V1 fusion timeseries from all other timeseries individually (for each subject), and calculated the Bayesian Evidence for a difference between the fusion timeseries. To test for category selectivity effects (place or person selective), we compared the fusion timeseries across categories within the ventral temporal and medial parietal ROIs separately. Here, we maintained the anatomical location (anterior and posterior) in our comparisons, which were pPPA-FFA1, aPPA-FFA2, Places1-Peope1, and Places2-People2. To compare the fusion timeseries across ventral temporal and medial parietal ROIs, we subtracted the medial parietal timeseries from their analogue in the ventral temporal cortex. We maintained the anatomical (anterior and posterior) and categorical (place or person selective) differences in our pairings, which were pPPA-Places1, aPPA-Places2, FFA1-People1, FFA2-People2.

## 3. Results

### 3.1 Vividness scores

We asked subjects to rate the average vividness of their mental imagery for each stimulus during each session (EEG, fMRI). This was to assess the range in vividness scores across subjects, and to identify any differences across stimulus categories or sessions. For each subject, we computed the average overall vividness score for the imagery of each stimulus during each session. During the EEG session, the mean vividness for people was 7.08 out of a maximum of 10 (SD = 1.11) and 6.74 for places (SD = 1.52). There was no significant difference between categories, t(11) = 0.881, p = 0.40. During the MRI session, the mean vividness for people was 6.88 out of 10 (SD = 1.13) and 6.67 for places (SD = 1.36). Again, there was no significant difference between categories, t(11) = 0.453, p = 0.659. The overall average vividness score was 6.91 for the EEG session and 6.77 for the MRI session. There was no significant difference between sessions, t(11) = 0.828, p = 0.425. The VVIQ scores ranged from 38 to 74, with an average score of 58.58.

### 3.2 MRI: whole-brain univariate contrast

First, we ran a whole-brain linear mixed model to quantify the univariate contrast of people versus places during mental imagery **(Figure 2)**. Regions with greater activity during the imagery of places included Places1 and Places2 in the medial parietal cortex, the lateral and ventral place memory areas (L/VPMA), the intraparietal sulcus (IPS), frontal eye fields (FEF), and a region in the inferior temporal sulcus (ITS) (all identified in both hemispheres). Greater activity during the imagery of people was found in People1 and People2 in the medial parietal cortex, the fusiform face area (FFA) (all identified in both hemispheres), and a small region in the right posterior superior temporal sulcus (pSTS). See the supplementary material for a full table of group level whole-brain results (S11). We also found a main effect of run in the peripheral visual cortex, where activity decreased across time, but no voxels where the interaction between run and category was significant.

**Figure 2.**
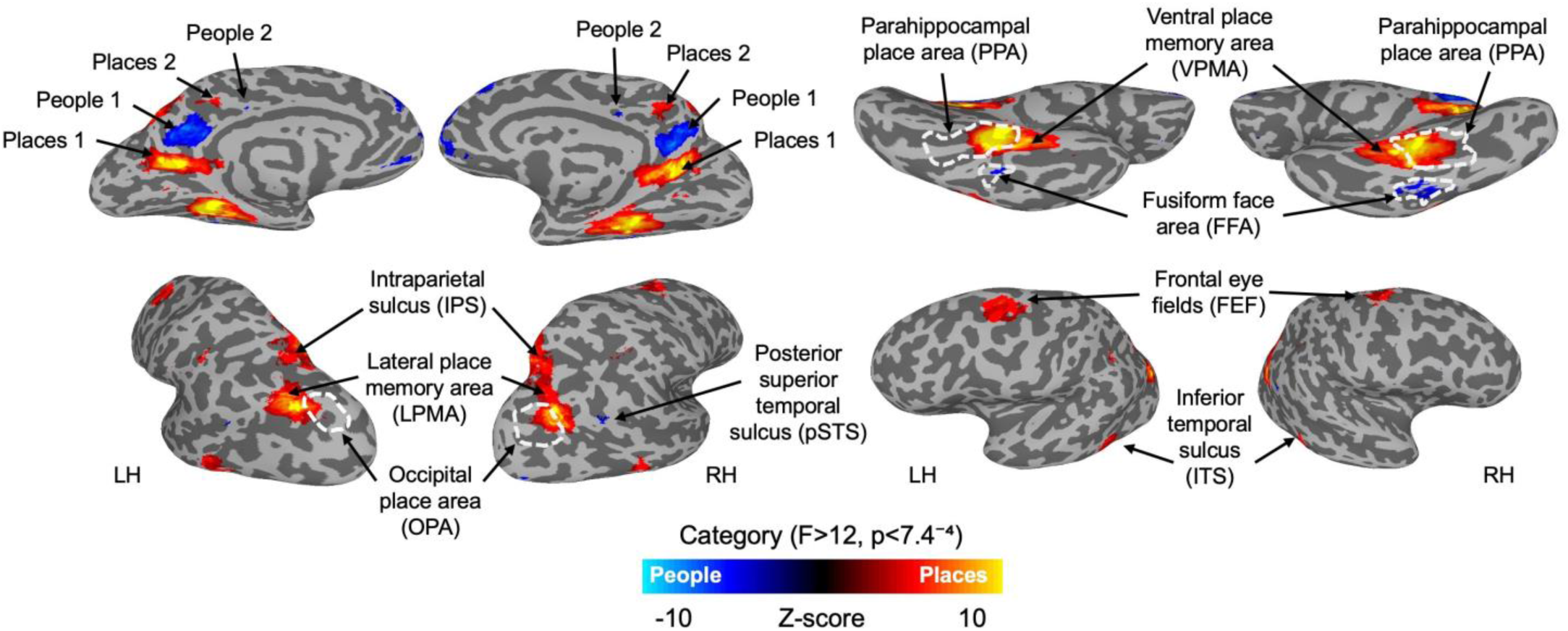
Group level results from the whole-brain contrast for the imagery of place versus people stimuli. The results are displayed on partially-inflated cortical surfaces of a sample subject, separately for the left and right hemispheres. Activation was thresholded at p < 7.4^-4^ (q < .0094) for the main effect of category, corresponding to a minimum F value of 12. Regions with greater activation for places are displayed in hot colours, and those with greater activation for people displayed in cold colours. Additional regions outlined in white dashed lines correspond to the group level activation from the perceptual scene/face localiser scans, contrasting perception of places versus people (t > 3.5, p < 0.0001 uncorrected). All regions were identifiable in both hemispheres.

The locations of our group level results in the medial parietal cortex were largely consistent with the locations of the ROIs taken from Silson et al. (2019), with the exception of lhPeople2, as indicated by their dice-coefficient scores (LH: Places1 = 0.85, People1 = 0.86, Places2 = 0.45, People2 = 0.02; RH: Places1 = 0.84, People1 = 0.80, Places2 = 0.59, People2 = 0.35). The small overlap for lhPeople2 is largely driven by its smaller size at the group level.

Also replicating previous findings, we found anterior shifts in the location of the lateral and ventral place memory areas in relation to their perceptual counterparts; the occipital place area (OPA) and PPA, respectively (Bainbridge et al., 2021; Steel et al., 2021). We found a smaller cluster of activation in the fusiform face area for imagery compared to the scene/face localiser scans, but unlike for places, perception and imagery of people activated very similar cortical locations. This mirrors recent findings reporting a lack of consistent shift in location for imagery of people (Chen et al., 2024).

In addition, we found a small region in the right posterior superior temporal sulcus (pSTS) with greater activation during imagery of people, consistent with previous reports (Kidder et al., 2025). It is worth noting that the cluster sizes for pSTS, Places2, People2, and FFA are small and would probably not survive a standard cluster-correction. Their size in the group level output would likely increase with the use of spatial soothing, but we opted against smoothing or further manipulating the data in this instance.

### 3.3 MRI: region of interest univariate analysis

Next, we extracted the univariate GLM beta values for people and place stimuli across medial parietal and ventral temporal ROIs. The medial parietal ROIs were taken from a prior dataset (Silson et al., 2019), and the ventral temporal regions were identified using independent scene/face localiser scans acquired on the same day as the imagery **(see Methods)**. We entered the average beta values (across hemispheres) within each ROI for each subject into a linear mixed model with factors Region and Category (people and places), keeping the medial parietal (People1, People2, Places1, Places2) and ventral temporal (FFA1, FFA2, pPPA, aPPA) regions separate. Across both sets of ROIs, activation was highest for the preferred category. In the medial parietal regions, there was a significant main effect of ROI (F(3,77) = 10.24, p = 9.50e-06), and an interaction between Category and ROI (F(1,77) = 26.89, p = 5.38e-12). The main effect of Category was not significant (F(1,77) = 2.71, p = 0.10). In the ventral temporal regions, there was a significant main effect of ROI (F(3,75.20) = 19.99, p =1.26e-9), a main effect of Category (F(1,75.06) = 8.64, p = 0.004), and an interaction between Category and ROI (F(1,75.06) = 24.27, p = 4.41e-11). Beta values plotted separately for each category within each ROI can be found in the supplementary material (S5). Univariate analysis including Hemisphere as a factor can be found in the supplementary material (S5). In summary, there was a main effect of Hemisphere in the medial parietal ROIs only, with slightly higher betas in the left hemisphere.

As a post-hoc analysis, we also tested whether the contrast values for places versus people stimuli during imagery were significantly different from zero in medial parietal and ventral temporal ROIs (Figure 3). This contrast value is simply the difference in activation across categories (i.e. the average beta for places minus the average beta for people in each ROI). The contrast was significantly greater than zero in Places 1, Places 2, aPPA, and pPPA ROIs, suggesting higher activation for places. The opposite was found in People 1, People 2, FFA1, and FFA2 ROIs, suggesting higher activation during the imagery of people (results from frequentist and Bayesian t-tests can be found in supplemental material S4). Thus, all of our ventral temporal and medial parietal ROIs demonstrated category selectivity during imagery of familiar stimuli, with greater activation for their preferred category. In comparison, in V1 the contrast values for places versus people were not significantly different from zero, suggesting that V1 was not category selective (t(11) = 0.969, p = 0.177). Contrast values plotted separately for each hemisphere can be found in the supplementary material (S5).

**Figure 3.**
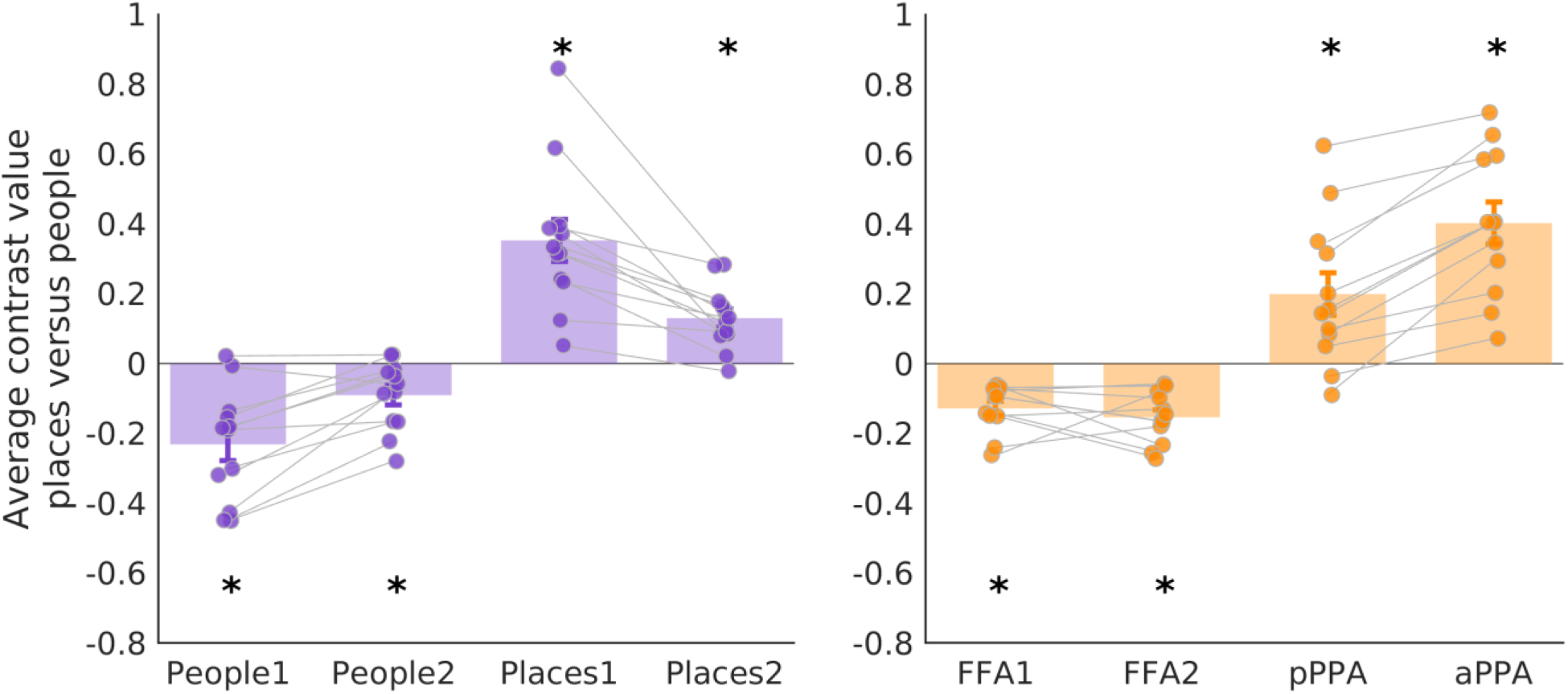
Average fMRI contrast values for places versus people imagery in the medial parietal (left panel) and ventral temporal (right panel), averaged across hemispheres. All contrast values were significantly different from zero, with values above zero in place selective regions and below zero in face selective regions (*p-values < FDR threshold: MPC p < .008, VTC p < .007). Each data point represents a single subject and error bars represent the standard error. FFA1 = posterior fusiform face area, FFA2 = anterior, pPPA = posterior parahippocampal place area, aPPA = anterior.

### 3.4 MRI: region of interest multivariate decoding

Our univariate ROI analysis confirmed that category selective regions in the medial parietal and ventral temporal cortex had higher response amplitudes during the imagery of their preferred category. However, differences in amplitude are not necessarily informative about the information content of these regions. First, we used multivariate pattern analysis to test whether we could also successfully decode the category of imagined stimuli across our ROIs (person or place). As expected, we were able to decode the category significantly above chance (50%) in all of our ROIs (∼60-70% accuracy) (see supplemental material S7 for Frequentist and Bayesian t-test results).

Next, we asked whether we could successfully decode the individual stimuli from each other within each category. We entered the average decoding (across hemispheres) within each ROI into two linear mixed models with factors Region and Category, keeping the medial parietal and ventral temporal regions separate for consistency with the statistical approach above. Our prediction was that we should be able to decode people stimuli and place stimuli from people- and place-selective regions, respectively.

In line with this prediction, in the ventral temporal regions there was only a significant interaction between Category and ROI (F(1,75.05) = 6.38, p = 0.0006), as decoding was found for each region’s preferred category, with the exception of FFA1 (see **Figure 4**). The main effect of ROI was not significant (F(3,75.42) = 0.45, p = 0.72), and the main effect of category was not significant (F(1,75.05) = 2.74, p = 0.10). In the medial parietal regions, the interaction between Category and ROI was not significant (F(3,77) = 1.04, p = 0.38). Instead, we found significant main effects of ROI (F(3,77) = 3.05, p = 0.03), and Category (F(1,77) = 7.06, p = 0.009). In Places2, we could decode the preferred category of place stimuli. However, in People1, People2, and Places2, we could decode stimuli in the non-preferred category only. Therefore, despite the clear category preferences in the medial parietal ROIs (indexed by differences in BOLD amplitude), the representation of different stimuli within these categories may be more complex (results from one-way t-tests against chance level decoding can be found in supplemental material S7). The ability to decode stimuli across different categories suggests that there may be information pertaining to both types of stimuli within these ROIs during imagery, which requires further investigation (see Discussion). Multivariate analysis including Hemisphere as a factor can be found in the supplementary material (S8). In summary, there were no significant main effects of Hemisphere in either medial parietal or ventral temporal cortex ROIs.

**Figure 4.**
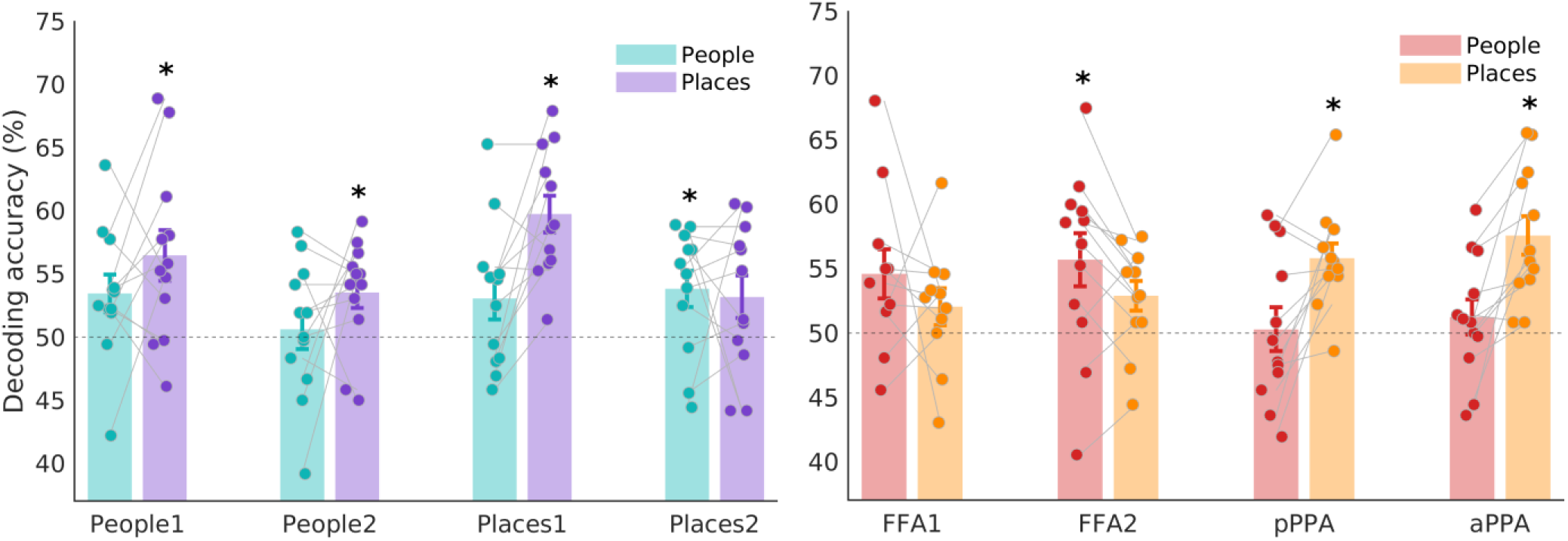
Average fMRI multivariate decoding of individual people and place stimuli in the medial parietal (left panel) and ventral temporal (right panel) regions of interest, averaged over hemispheres. Chance level decoding was 50% as we took the average decoding across all pairs of stimuli (indicated by the dashed line). Asterisks indicate stimulus decoding significantly above chance accuracy (*p-values < FDR threshold: MPC p < .022, VTC p < .019). Each data point represents a single subject and error bars represent the standard error. FFA1 = posterior fusiform face area, FFA2 = anterior, pPPA = posterior parahippocampal place area, aPPA = anterior.

Next, we checked stimulus decoding in V1. The perception of people and places has been attributed to processing foveal versus peripheral information, respectively (Levy et al., 2001). During perception, face-selective regions have a more pronounced foveal bias, whereas scene-selective regions are biased towards the periphery (Silson et al., 2015). Given the typical association between category and eccentricity, we also asked whether it was possible to decode individual stimuli within foveal and peripheral portions of V1 during visual imagery. We considered whether people stimuli would be more easily decoded in foveal V1, whereas place stimuli would be more easily decoded in peripheral V1. Contrary to these predictions, we found significant decoding of places in both portions of V1, but no significant decoding of people (see Figure 5).

**Figure 5.**
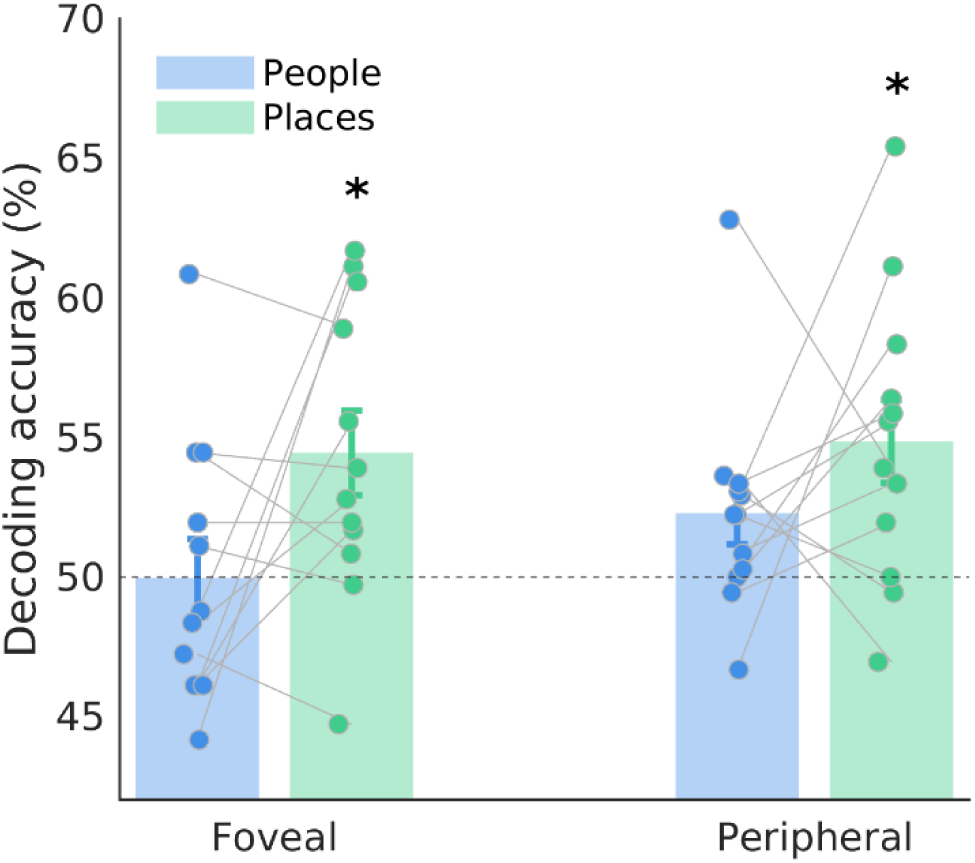
Average fMRI multivariate decoding of individual people and place stimuli in the foveal and peripheral divisions of V1, averaged across hemispheres. Chance level decoding is 50% (indicated by the dashed line). Asterisks indicate decoding that was significantly above chance accuracy (*p-values < .013 FDR threshold). There was significant decoding of places across both regions of V1, but no above chance decoding of people. Each data point represents a single subject.

We hypothesise that this may be due to the differences in the visual properties of the people and place stimuli subjects were cued to imagine, as there is generally more variation across places than people. To test this assumption, we ran the images provided to us by our subjects through two image computable models (gist and LGN) and constructed an RDM across these images for each subject (using Euclidean distance). We then calculated the average dissimilarity across people and place images separately for each subject, and ran a paired t-test across these dissimilarities. There was significantly higher dissimilarity within the place stimuli compared to people using the LGN model (t(11) = 3.03, p = .011), but no significant difference using the gist model (t(11) = 1.37, p = .197), although dissimilarity was numerically larger for places compared to people. This difference likely reflects the different ways that these models spatially sample the images.

To summarise the fMRI results, we successfully identified people and place memory regions within medial parietal cortex in a whole brain univariate analysis (Silson et al., 2019), and confirmed that activation within these regions was greatest during imagery of their preferred category. We also demonstrated that regions within ventral temporal cortex that are typically considered visual (FFA1/2 and p/aPPA) were also active during imagery, and had the expected category selectivity. Using multivariate decoding analysis, we tested the hypothesis that we would be able to successfully decode individual stimuli within the preferred category for each ROI. This was largely the case in the ventral temporal ROIs (with the exception of FFA1, where decoding of people stimuli did not pass the FDR post-hoc correction for multiple comparisons), but a more complex pattern emerged in the medial parietal cortex; these regions represented information during imagery that was not restricted to their preferred category. In comparison, we were only able to decode place stimuli in foveal and peripheral V1 (with no above chance decoding of people stimuli), which we hypothesise may be due to the increased visual variability in the to-be-visualised place stimuli compared to the people. Overall, our MRI results confirmed the roles of our key ROIs in visual mental imagery, and hint that there might be shifts in the information content across medial parietal, ventral temporal, and primary visual ROIs. Next, we used the EEG data to investigate the timecourse of this imagery information, and identify timepoints where we could successfully decode both the category and identity of mental images across the imagery period.

### 3.5 EEG: multivariate decoding through time

Multivariate pattern analysis (MVPA) was first used to identify timepoints where the category of personally familiar stimuli (people or places) could be decoded above chance accuracy across all EEG electrodes. Stimulus category was highly decodable for the first 700ms of the imagery period, suggesting an early difference in the activity associated with mental imagery for these categories **(Figure 6, top panel)**. This early higher decoding was followed by sustained but weaker category information with a lower accuracy and variable Bayesian evidence. We speculate that this may reflect differences in the prolonged maintenance of mental images across participants, stimuli, and individual trials. Next, we took the average decoding within each category (6 stimuli in each category) **(Figure 6, bottom left and right panels)**. We present the average across all comparisons so that all plots have chance level decoding of 50%. Here, individually imagined stimuli were decodable from each other for a shorter time period following imagery onset, lasting roughly 100ms. However, there was a second peak in decoding for both categories of stimuli at ∼700ms into the imagery period (∼1200ms from cue onset) for places and ∼1100ms for people (∼1600ms from cue onset).

**Figure 6.**
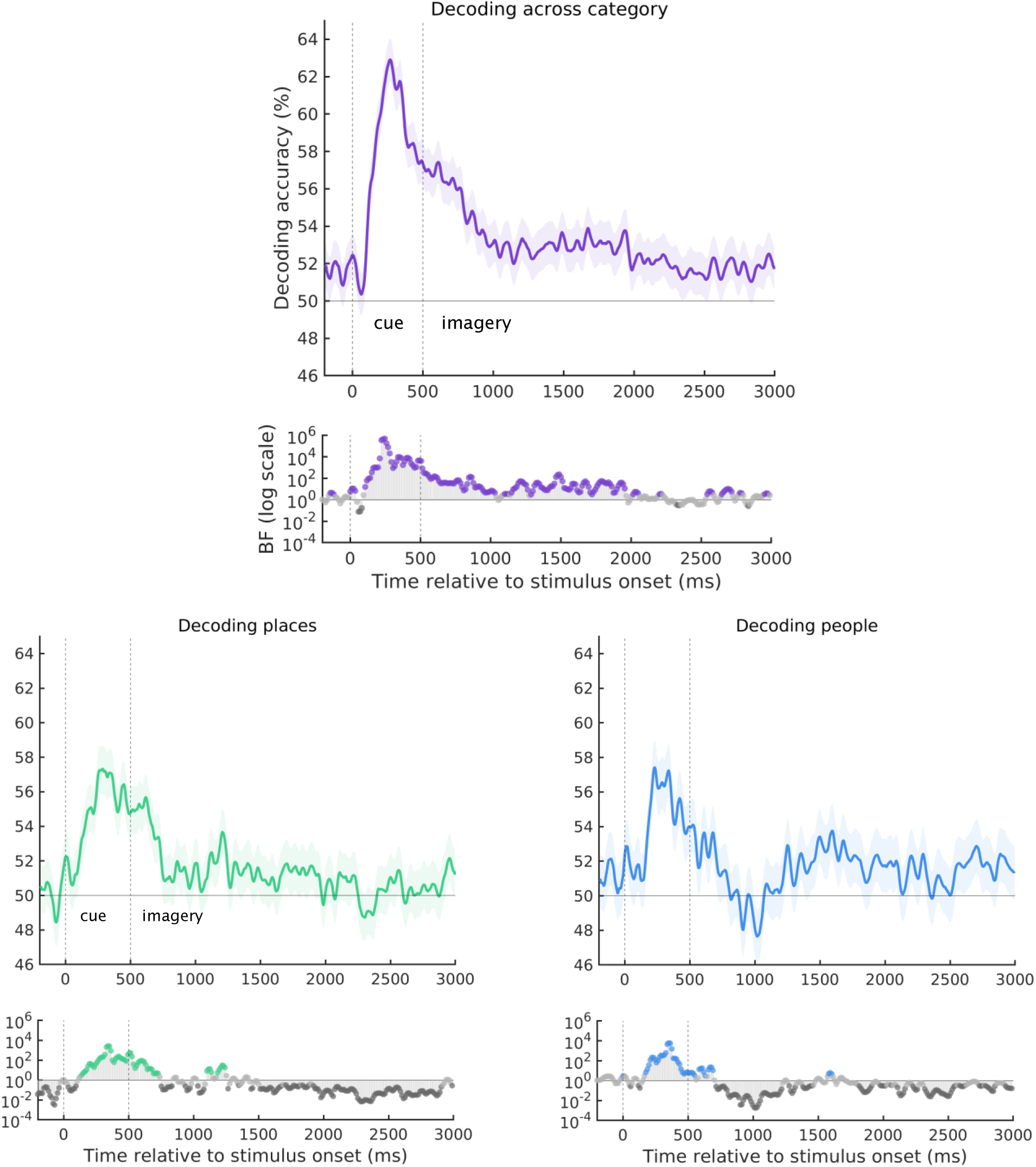
Average multivariate EEG timeseries decoding during imagery. **Top:** across category decoding of people versus place stimuli. **Bottom left:** within category decoding of places. **Bottom right:** within category decoding of people. The cue was presented during the first 500ms, followed by 2500ms of imagery. Here we plot both average decoding across subjects and the standard error around the mean, smoothed with a moving average of 10 samples for visualisation. Chance level decoding was 50% for all analyses as we took the average decoding across all pairs of stimuli. Timepoints with Bayes Factors that were greater than 3 are highlighted by the coloured points, suggesting that the alternative hypothesis of decoding above chance level was at least three times more likely than the null hypothesis (BF_10_ >3). Dark grey points indicate Bayes Evidence in favour of the null hypothesis (BF_10_ <1/3).

We then used time-generalised decoding to examine the stability of imagery representations across time. Increased decoding along the diagonal indicates that the classifier can only generalise to its neighbouring timepoints and that activity is changing over time. In comparison, off-diagonal decoding suggests sustained representations that are more stable through time (King et al., 2014). Here category decoding (people or places) throughout the trial successfully generalised to both an early (∼600-1000ms) and late test window (∼1600-2100ms), indicating two peaks of sustained imagery representation throughout the trial **(Figure 7)**. Importantly, there was little evidence for the cue period time-generalising to the imagery period, reassuring us that category decoding was not driven solely by the presence of the cue.

**Figure 7.**
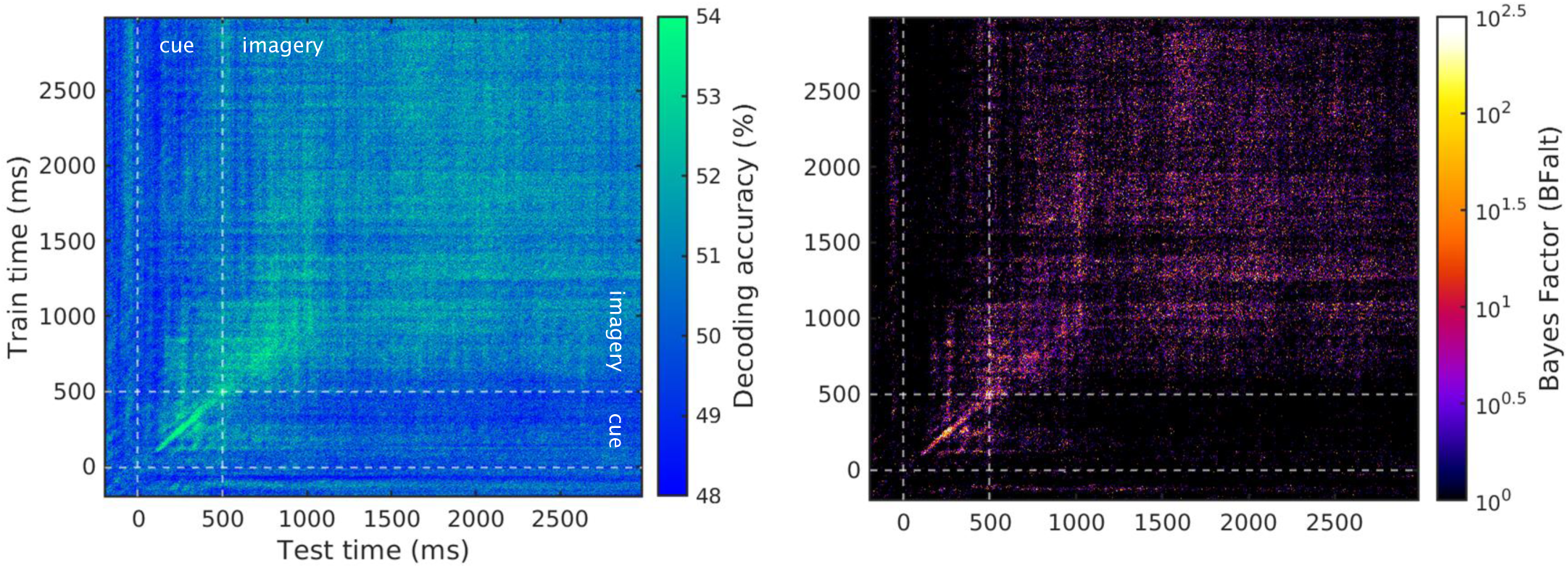
Average time-generalised EEG decoding during imagery. **Left:** Category level time-generalised decoding of people versus place stimuli across all timepoints within the imagery period. The cue was presented during the first 500ms, followed by 2500ms of imagery. **Right:** Bayes Factors indicating the evidence for the alternative hypothesis, plotted on a log10 scale. Brighter colours indicate decoding above chance level (50%).

To summarise the EEG analysis, we were able to successfully decode category (places versus people) throughout the imagery period. Decoding of within-category stimuli was weaker, possibly due to the reduced number of trials available for classification and the subtler differences between stimuli. However, we were still able to decode both people and place stimuli during two distinct time windows. Stimuli in both categories were decodable during the first ∼100ms of the imagery period, followed by a re-emergence of decodable activation at ∼700ms into the imagery period for places and ∼1100ms for people. Time-generalised decoding indicated that the imagery was sustained throughout the imagery period, with relatively stable representations that appeared to form early and late clusters. These results confirmed that information pertaining to both the category and stimulus identity was captured in the EEG data, but provide no spatial information. Next, we combined the EEG and fMRI data to identify when in time the representational structures present in the fMRI ROIs matched those captured in the EEG.

### 3.6 EEG and fMRI fusion

Lastly, we calculated the EEG-fMRI fusion time series by correlating the fMRI RDMs in each ROI with the EEG RDMs at each time point. As shown in **Figure 8A**, all regions had at least some timepoints with strong Bayesian evidence for a correlation between the EEG and fMRI RDMs (although the evidence for fusion in People2 and FFA2 was weaker). In the medial parietal ROIs, evidence for clusters of significant fusion (BF_10_ > 3) occurred first in Places2 (500ms) and Places1 (539ms), followed by People1 (691ms). In People2, there was only a small window with significant fusion (at 852ms). In V1, sustained fusion started later at 1371ms (with a small window at 984ms lasting only 12ms). This fusion pattern may indicate a feedback sweep from place selective regions within the medial parietal cortex to V1.

**Figure 8.**
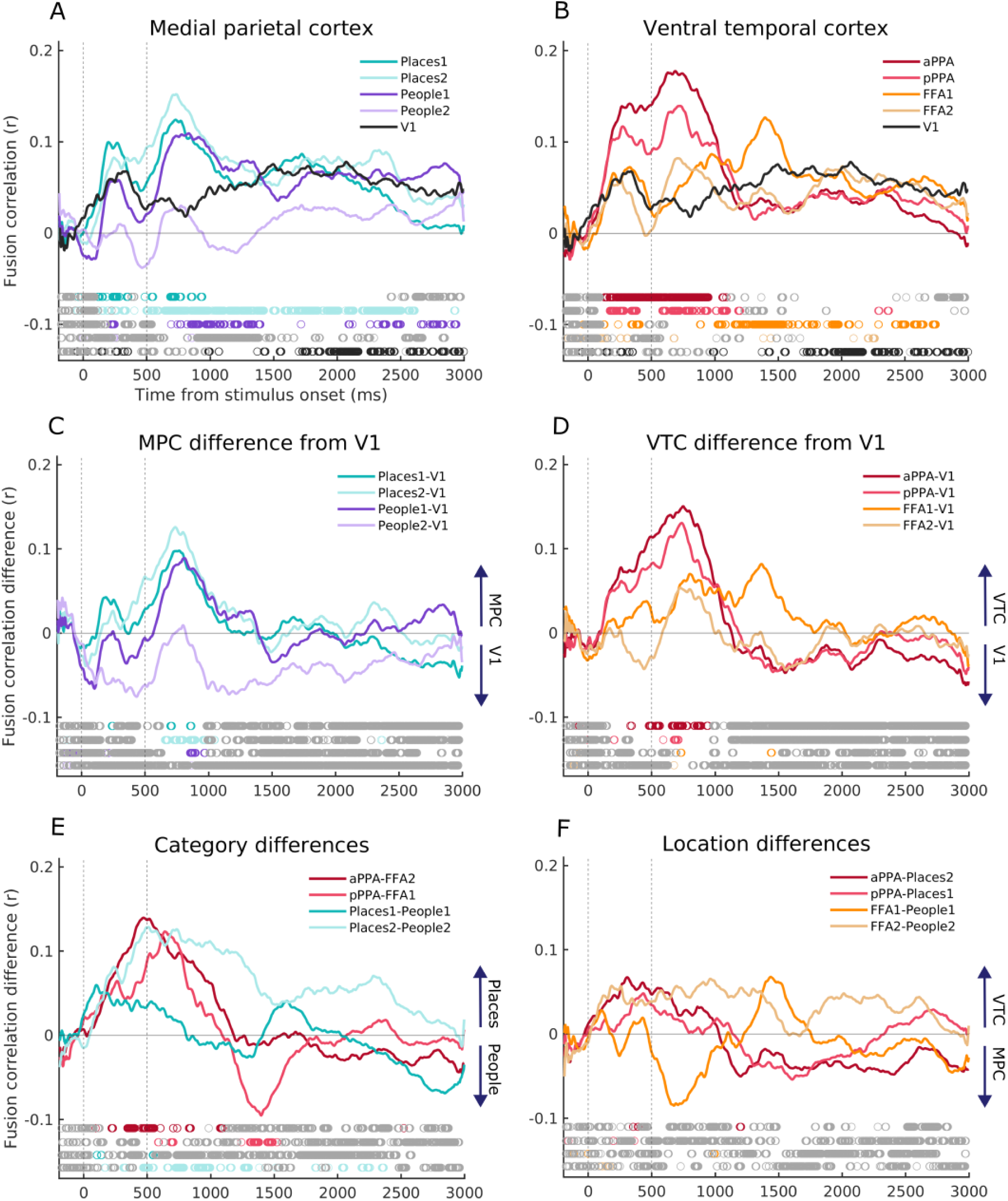
**Top:** Average EEG-fMRI fusion timeseries for the medial parietal (**A**) and ventral temporal (**B**) regions of interest, averaged over hemispheres. The V1 timeseries (plotted in black) is replicated across plots A and B to aid comparison across ROIs. **Middle:** The difference between the fusion timeseries of medial parietal (**C**) and ventral temporal (**D**) ROIs and V1 (where the V1 fusion timeseries was subtracted from the other ROIs). Positive values indicate that the ROI had a higher fusion correlation than in V1. **Bottom:** (**E**) The difference between fusion timeseries in place selective ROIs compared to their face selective counterparts (Category differences). Positive values indicate that place selective ROIs had a higher fusion correlation than their people selective counterpart, while negative values indicate higher fusion in people selective ROIs. (**F**) The difference between fusion timeseries across ventral temporal cortex and medial parietal cortex (Location differences), maintaining their anatomical (anterior or posterior) and categorical relationships (people or place selective). Positive values indicate a higher fusion correlation in ventral temporal cortex ROIs compared to their medial parietal counterparts, while negative values indicate a higher fusion correlation in medial parietal ROIs. **All:** Timepoints with Bayes Factors that were greater than 3 are indicated by the coloured circles along the bottom bar, suggesting that the alternative hypothesis of a fusion correlation is at least three times more likely than the null hypothesis (BF_10_ >3). Grey circles indicate Bayes Evidence in favour of the null hypothesis (BF_10_ < 1/3).

In the ventral temporal ROIs, a similar pattern emerged (**Figure 8B**). Fusion arose first in aPPA (500ms), followed by pPPA (586ms), and FFA1 (883ms). FFA2 had a few small windows with BF_10_ > 3 (676ms, 1383ms, and 2207ms), but little evidence for any sustained fusion over time. In general, this finding mirrors the result above with fusion peaking first in place selective regions, followed by face selective regions, and then early visual cortex.

To directly compare the fusion correlation timings across ROIs and conditions, we calculated a series of difference waves. First, to identify differences between the V1 fusion and other ROIs, we subtracted the V1 fusion timeseries from all other timeseries individually (for each subject), and calculated the Bayesian Evidence for a difference between the fusion timeseries (**Figure 8C and D**). There was Bayesian evidence (BF_10_ > 3) for higher fusion correlations compared to V1 in most ROIs during the early imagery time window (up to ∼500ms post imagery onset). This confirms that there was a difference in fusion correlation between our category-selective ROIs and V1 during the first 500ms of imagery (with the exception of People2 and FFA2, which had lower fusion correlations in general). Subtracting the V1 fusion timeseries also reduced the fusion correlations during the cue period, where the word cue was present on the screen and the fusion is likely to be driven by the visual input to all ROIs.

In addition, we compared the fusion timeseries across categories within the ventral temporal and medial parietal ROIs separately. Here, we maintained the anatomical location (anterior and posterior) in our comparisons, which were pPPA-FFA1, aPPA-FFA2, Places1-Peope1, and Places2-People2. In ventral temporal cortex (**Figure 8E**), we found Bayesian evidence (BF_10_ > 3) for higher fusion in place selective regions (a/pPPA) compared to face selective regions (FFA1/2) during the early imagery window (up to ∼500ms into imagery). At a later timepoint, we also found evidence of the reverse, with higher fusion in FFA2 compared to aPPA between 800-1000ms into the imagery period. Therefore, in the ventral temporal cortex, fusion was higher in place selective regions during an early time window, and higher in face selective regions in a later time window. The difference across categories in medial parietal cortex was less clear (**Figure 8E**), with greater fusion across the whole imagery window in Places2 compared to People2. However, this is not surprising given that there was little evidence for any fusion in People2 at all, and this difference is not necessarily category specific. In comparison, the Bayesian evidence for the difference between Places1 and People1 was predominantly in favour of the null hypothesis (BF_10_ < 1/3), indicating similar fusion time courses. Therefore, although there appear to be small differences in the peak timing of fusion across categories within medial parietal cortex, the evidence in support of this is mixed.

Next, to compare the fusion timeseries across ventral temporal and medial parietal ROIs, we subtracted the medial parietal timeseries from their analogue in the ventral temporal cortex (**Figure 8F**). We maintained the anatomical (anterior and posterior) and categorical (place or person selective) differences in our pairings, which were pPPA-Places1, aPPA-Places2, FFA1-People1, FFA2-People2. We found Bayesian evidence predominantly in favour of the null hypothesis (BF_10_ < 1/3), indicating no difference in the fusion timeseries across these ROI pairs.

As some meta-analyses have indicated lateralised effects in perceptual and memory regions (Spagna et al., 2021; Winlove et al., 2018), we also compared the fusion across left and right hemisphere ROIs. Here, we found Bayesian evidence predominantly in favour of the null hypothesis, indicating no difference in the fusion timeseries across hemispheres (see supplementary material S9).

## 4. Discussion

Here we used an EEG-fMRI fusion approach to investigate the involvement of medial parietal and ventral temporal category-selective regions during mental imagery. We also compared the activity within these higher-level regions to that in V1. We hypothesised that we may find differences in the fusion time courses for medial parietal and ventral temporal ROIs during mental imagery, assuming that they may play distinct roles. If mental imagery is supported by a mechanism of perception in reverse, then we may expect to see activity peak earlier in medial parietal cortex than ventral temporal given both their anatomical locations on the cortex (MPC regions are more anterior) and the prominent role that VTC regions play in visual perception. However, we did not find any clear differences in the fusion timecourse across these ROIs, with Bayesian evidence predominantly in favour of the null hypothesis. This temporal similarity between two spatially distinct cortical regions mirrors recent electrocorticography (ECoG) data during recognition of famous people and places, where medial parietal and medial temporal regions had similar time courses of activation (Woolnough et al., 2020). One possible explanation for this is that both sets of regions received input from elsewhere, driving all regions to represent visual imagery simultaneously (Spagna et al., 2024). Candidate regions include those identified in our whole-brain univariate analysis, for example the lateral place memory area. Further, the recruitment of subcortical brain structures such as the hippocampus and amygdala are known to support memory and associated mental imagery processes (Argyropoulos et al., 2019; Ranganath & Ritchey, 2012; Ritchey & Cooper, 2020; Silson, Steel, et al., 2019; Spagna et al., 2024; Treder et al., 2021), and could also be driving the activity across our main ROIs. Prior work has suggested that the hippocampus (in particular the anterior portion, anterior PPA), and portions of the medial parietal cortex form a distinct scene memory network that links visual processing with long-term memory and additional scene context (Baldassano et al., 2016; Tullo et al., 2023). Here we replicated findings of category-selectivity in the hippocampus (reported in supplementary material S6), which had higher activity during the imagery of place stimuli compared to people.

Instead of a difference in fusion correlation driven by ROI location, we found a difference driven by the category; in both cases, regions selective for places (Places1/2, p/aPPA) had a higher correlation earlier in time than those selective for people (People1/2, FFA1/2). It is possible that, in this sample of subjects, mental imagery of place stimuli was quicker to emerge than people. Although we did not formally assess this, some subjects did report that place stimuli took less time to visualise in their end of session questionnaires, despite there being no overall difference in reported subjective vividness. Future behavioural investigation is required to establish if this is a consistent finding and the extent of any between-subject variation.

Using multivariate pattern analysis, we were able to significantly decode not just category-level information, but also the individual within-category stimuli being visualised during the imagery period. We predicted that we would be able to decode imagery of individual people in regions that are ‘selective’ for face stimuli (i.e. FFA1/2 and People1/2) in our fMRI data. Similarly, we expected to decode individual place stimuli in regions selective for place stimuli (i.e. p/aPPA and Places1/2). This prediction was confirmed in the ventral temporal ROIs, where we found a significant interaction between ROI and category. Post-hoc tests for significant decoding within each ROI identified decoding that was confined to the preferred stimulus category. The contradiction to our hypothesis was more pronounced in the medial parietal regions, where we only found a main effect of category with no significant ROI interaction. While there was clear category-selectivity in the mean univariate activation for people and place selective ROIs, the information content within medial parietal ROIs was not solely related to their preferred category during mental imagery.

Our fMRI ROI stimulus decoding, together with the lack of clear differences in the fusion onset timings across medial parietal and ventral temporal cortex, leaves an open question about what information is represented across high-level medial parietal ROIs during mental imagery. Although the patterns of stimulus decoding differed across these regions, the representational structures correlated with the EEG at similar points in time to the ventral temporal ROIs. Stimulus category was not the only difference between the stimuli provided by our subjects; other dimensions such as familiarity, emotional relevance, semantic associations etc. may also structure the representations during mental imagery within medial parietal cortex specifically. Indeed, similar regions in medial parietal cortex have been associated with the semantic structure of people and place images (Koslov et al., 2024) and varying spatial scales (Peer et al., 2019) to name only a few (for a review see Foster et al., 2023). Future work could focus on sampling a larger range of stimuli across other possible dimensions and evaluate how this links to stimulus decoding in medial parietal cortex.

Several studies suggest a left hemisphere dominance during mental imagery, particularly in the left temporal lobe (Bartolomeo, 2002, 2008; Moro et al., 2008; Thorudottir et al., 2020), and a ventral temporal region known as the Fusiform Imagery Node (Hajhajate et al., 2022; Spagna et al., 2021). This region sits in cytoarchitectonic sector FG4 of the fusiform-gyrus (Lorenz et al., 2017), and is likely to overlap with FFA2. Other work has also suggested left-lateralised effects in the medial parietal cortex during scene imagery, corresponding roughly to Places1 and People1 in our analyses (Silson, Gilmore, et al., 2019). We found only one significant main effect of hemisphere across our analyses, with significantly greater univariate activation of the left hemisphere medial parietal regions compared to the right. This was not found for the ventral temporal ROIs, and the effects of hemisphere in our decoding analyses were also not significant for either medial parietal or ventral temporal regions. Further, the Bayesian evidence for the comparison between left and right hemisphere ROIs in the fusion analyses indicated evidence predominantly in favour of the null hypothesis. Therefore, we found only limited evidence for lateralised effects in the present data. However, our analyses differ from other studies in that we predominantly compared the imagery of people versus places, while lateralised effects are often reported for the comparison between imagery and rest, or some other control condition (Spagna et al., 2021; Winlove et al., 2018). The lateralisation of mental imagery may also be dependent on the specific item to be visualised (Liu et al., 2022). In addition, we recruited both left and right handed subjects, which may complicate the effects across the group as found for other perceptual regions (Thome et al., 2022).

Using EEG-fMRI fusion, we found that the representations of people and place imagery in V1 correlated with EEG signals at a later timepoint than either high-level category-selective regions in medial parietal or ventral temporal cortex. This suggests that the activity in V1 may be related to feedback or recurrent processing that emerged later in time than medial parietal and ventral temporal ROIs. Does this finding mean that imagery was visual in nature? Using multivariate pattern analysis in the fMRI data, we were able to decode the individual place stimuli during imagery in both central and peripheral V1. This indicates that there were quantifiably distinct patterns of activation associated with imagery of each place in V1, which could suggest ‘perception-like’ imagery and the mechanism of perception in reverse. However, the classifier could use any number of features to distinguish between stimuli, and decodable signals within V1 could reflect any number of processes that are not necessarily identical to visual perception (Favila et al., 2022; Naselaris et al., 2015; Pace et al., 2023).

Unlike the place stimuli, we were not able to significantly decode people stimuli, either in central or peripheral V1. One possible explanation for this is that the patterns in V1 during imagery of places were more varied in structure than for people, resulting in better decoding for these stimuli. Indeed, the visual distinctiveness of the place stimuli was greater than that of the people when we ran the subjects’ images through an LGN visual model, which may explain the differences in V1 decoding across categories. However, this explanation makes the explicit assumption that imagery activity in V1 is ‘perception-like’, and that subjects were faithfully recreating the images that they provided during imagery (which may not have always been the case).

Another question is whether V1 activity is *necessary* for mental imagery (Spagna et al., 2024). Patient findings indicate that mental imagery is possible despite visual cortex damage, strongly suggesting that V1 is not a critical region in supporting imagery (Della Sala & Spinnler, 1988; Hajhajate et al., 2022). Activity in V1 may also not be *sufficient* for our conscious experience of imagery, as sensory information can be decoded in V1 in Aphantasic individuals who lack visual imagery (Cabbai et al., 2024). Given the nature of our analysis and the timing of the activity in V1, we do not make any claims about causality; our results simply highlight differences in the timing of stimuli representations in medial parietal, ventral temporal, and primary visual cortex during mental imagery in healthy individuals.

One limitation of this study is the low sample size, driven by the constraints of running a multisession within-subject neuroimaging study. We also acknowledge that our decoding accuracies for the decoding of individual stimuli ranged from 54 to 60%. While this is lower than the decoding accuracies that you might achieve in studies of visual perception, we emphasise the difficulty of decoding individual stimuli during mental imagery (e.g. imagining mum versus dad). This analysis relies on the patterns of information across these within-category stimuli being sufficiently different to facilitate above chance decoding. When running the easier classification of category in our ROIs (i.e. people versus places), we are able to decode with higher accuracies ranging from 60 to 75%. Future studies could potentially improve stimulus level decoding by minimising the visual and semantic overlap between the familiar stimuli, and by training participants to generate similar mental images for each exemplar at each trial repeat.

In summary, we used EEG-fMRI fusion to investigate the time course of mental imagery representations within category-selective regions in medial parietal, ventral temporal cortex, and V1. The fusion correlations peaked early on during imagery and did not clearly differ across medial parietal and ventral temporal ROIs. However, during an early time window, fusion was higher in place selective regions, whereas fusion in face selective regions was higher during a later time window. This mirrors EEG stimulus decoding results where the peak decoding of individual places was earlier in time than peak decoding of people. In V1, fusion correlations occurred later during the imagery period, possibly reflecting the top-down progression of mental imagery from category selective regions to primary visual cortex. Within the ventral temporal ROIs, fMRI decoding was highest for the preferred category. In contrast, a more complex pattern of decoding was identified in the medial parietal cortex. These regions represented information during imagery that was not restricted to their preferred category, prompting future investigation to establish how these regions support the generation of mental images. Overall, our results confirm the engagement of category-selective cortex during visual mental imagery, and evidence shifts in information content across medial parietal, ventral temporal, and primary visual cortex during this complex and dynamic process. Understanding the extent to which medial parietal regions represent category information according to different dimensions (visual, semantic, emotional) remains an avenue for future research.

## Acknowledgements

This work was funded by the Biotechnology and Biological Sciences Research Council (BBSRC) BB/V003917/1, and the School of Philosophy, Psychology and Language Sciences at the University of Edinburgh (UoE). The MRI data was collected at the Edinburgh Imaging Facility. For the purpose of open access, the author has applied a creative commons attribution (CC BY) licence to any author accepted manuscript version arising.

## Data availability

Data are available on the Open Science Framework: https://osf.io/ypk6c/

## CRediT contributor roles

**CLS:** Conceptualisation, funding acquisition (UoE), investigation, data curation, formal analysis, visualisation, writing - original draft, writing - reviewing and editing. **JAT:** writing - review and editing, visualisation. **EHS:** Conceptualisation, funding acquisition (BBSRC and UoE), formal analysis, writing - review and editing, supervision.

## Notes

https://osf.io/ypk6c/

### Competing Interest Statement

The authors have declared no competing interest.

### Summary of Updates

Updated and clearer methods, figures, and supplementary material.

