## supplementary material for "Visual imagery of familiar people and places in category selective cortex"

**S1. Wording of the request for stimuli**

‘For the experiment, we would like you to provide us with 6 photos of people that are personally familiar to you, and 6 photos of places that are personally familiar to you. It does not matter exactly who/what these images are, only that you are familiar with what they look like, and you are confident that you could create a reasonable mental image of these people and places. For example, they could be photos of your best friend, your partner, your childhood home, your University flat, your favourite museum etc. The photos of people should be headshots, containing mostly their face and not their whole body. The place images should not contain any people. Please give the photos relevant names, such as 'Sophie’, or ‘Office’, as these are the descriptions that we will use when we ask you to recall the images during the experiment’.

**S2. Number of nodes used in the fMRI analysis**

**Ventral temporal cortex:** The PPA and FFA ROIs were drawn manually for each subject on the surface (SUMA draw ROI) after thresholding the contrast of scenes versus faces from the scene-face localiser at t > 3.5 (p<0.0001, uncorrected). PPA and FFA ROIs were divided into anterior and posterior portions, either by identifying two separate peaks in the t-maps, or by dividing a single large cluster equally along the posterior-anterior axis (Baldassano et al., 2016; Silson et al., 2019). Not all ROIs were identifiable across all subjects (left hemisphere: FFA1 = 10, FFA2 = 12, pPPA = 9, aPPA = 12; right hemisphere: FFA1 = 10, FFA2 = 11, pPPA = 12, aPPA = 12). The number of nodes included in each ROI for each subject are listed in the table below.

| **Subject** | **lhFFA1** | **lhFFA2** | **lhpPPA** | **lhaPPA** | **rhFFA1** | **rhFFA2** | **rhpPPA** | **rhaPPA** |
| --- | --- | --- | --- | --- | --- | --- | --- | --- |
| 1 | 673 | 443 | 826 | 2232 | NaN | 319 | 410 | 732 |
| 2 | 358 | 216 | 838 | 731 | 318 | 199 | 962 | 542 |
| 3 | 303 | 1214 | 798 | 959 | 374 | 322 | 757 | 859 |
| 4 | 261 | 146 | 910 | 1074 | 297 | 359 | 959 | 793 |
| 5 | 572 | 432 | 1015 | 1227 | 646 | NaN | 911 | 1197 |
| 6 | 297 | 321 | 1361 | 1683 | 508 | 455 | 593 | 925 |
| 7 | NaN | 380 | NaN | 912 | 345 | 360 | 558 | 551 |
| 8 | 201 | 190 | 1104 | 1278 | 372 | 285 | 1057 | 889 |
| 9 | 290 | 354 | NaN | 1090 | 328 | 314 | 765 | 610 |
| 10 | 673 | 472 | 764 | 854 | 305 | 429 | 1037 | 1099 |
| 11 | NaN | 348 | NaN | 1170 | NaN | 302 | 484 | 1015 |
| 12 | 613 | 681 | 715 | 820 | 765 | 273 | 941 | 840 |
| **mean** | 424 | 433 | 926 | 1169 | 426 | 329 | 786 | 838 |
| **SD** | 185.89 | 284.51 | 204.32 | 420.79 | 161.51 | 71.28 | 225.65 | 207.62 |

**Medial parietal cortex:** Medial parietal ROIs, also defined on the surface, were taken from previous work (Silson et al., 2019). Here, two ROIs on the medial parietal surface with increased BOLD activation during imagery of place stimuli were named as Places1 (left hemisphere = 2587 nodes, right hemisphere = 2766) and Places2 (LH = 651 nodes, RH = 1050). A further two ROIs with greater activation for people stimuli were named People1 (LH = 1319 nodes, RH = 1455) and People2 (LH = 136 nodes, RH = 215).

**S3.** **MRI: region of interest average timeseries correlations**

To visualise the similarity in fMRI timecourses across our main regions of interest, we created a group level dissimilarity matrix across all runs and subjects **(Figure S1)**. This provides a simple pictorial representation of the fMRI timeseries correlations during mental imagery across ventral temporal cortex, medial parietal cortex, and primary visual cortex. To do this, we extracted the mean of the pre-processed timeseries within each volume ROI in each hemisphere during each run of the MRI recall data (3dmaskave). For each subject, separate RDMs for each ROI in each hemisphere were constructed using the recall deconvolution where each stimulus was modelled with one regressor across all runs. The t-values estimated for each of the 12 stimuli across the ROI nodes extracted from the surface data were used to construct 12x12 RDMs (1 – Pearson’s correlation r values) ordered by their category (people then places). We then constructed a group level RDM by averaging across all subject-level RDMs.

Previous fMRI results indicate increased connectivity between aPPA and Places1, and FFA2 and People1 (Silson et al., 2019; Koslov et al., 2024; Baldassano et al., 2016). Here, we found that the aPPA fMRI timeseries during imagery was most similar to (or least dissimilar from) both pPPA and Places1, and most dissimilar to People1 and People2 **(Figure S1)**. These correlations were relatively stable across left and right hemispheres, and all ROIs had high correspondence with their homologue in the opposite hemisphere (as indicated by the diagonal of dark navy squares in the bottom left/top right quadrant). People1 and People2 were the most dissimilar to the other ROIs included here, but were closer in profile to FFA1 and FFA2, as expected. These differences in correlation across category selective regions likely reflect increased activation during the mental imagery of stimuli from their preferred category, which occurred at different timepoints within each run.

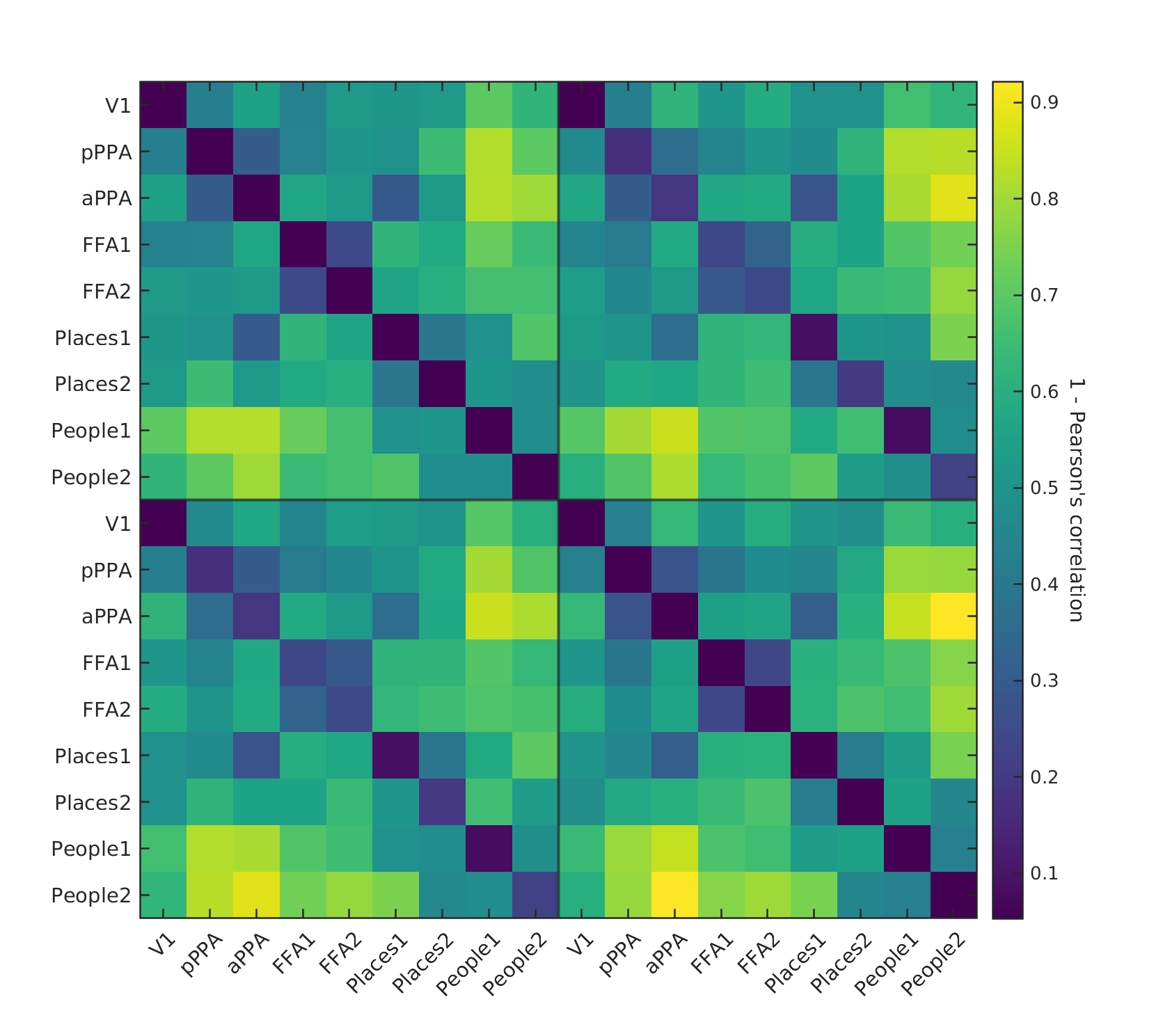

Left

Left

Right

Right

**Figure S1:** Group average representational dissimilarity matrix (1 - Pearson’s correlation) for the fMRI timeseries across regions of interest. Light green and yellow colours indicate the greatest dissimilarity in the timeseries, whereas dark green and navy indicate the least dissimilarity (or most similarity). RDMs were calculated within each run for each subject separately before being averaged to create a group level RDM.

**S4. MRI region of interest univariate analysis: MPC, VTC, and V1**

Results from one-sample Student and Bayesian t-tests for the contrast of places versus people in ventral temporal and medial parietal ROIs shown in **Figure 4.** We also provide the same values for V1. Regions with contrast values significantly greater than zero had increased activity for place imagery compared to people, and vice versa (FDR thresholds: MPC p < .008, VTC p < .007)

| **One Sample T-Test** | | | | | | | | | | | |
| --- | --- | --- | --- | --- | --- | --- | --- | --- | --- | --- | --- |
|  | | **t** | | **df** | | | **p** | | **Cohen's d** | | **SE Cohen's d** |
| **PPAp** |  | -3.276 |  | 11 | |  | 0.007 |  | -0.946 |  | 0.347 |
| **PPAa** |  | -6.783 |  | 11 | |  | 3.020×10^-5^ |  | -1.958 |  | 0.493 |
| **FFA1** |  | 6.124 |  | 10 | |  | 1.121×10^-4^ |  | 1.846 |  | 0.496 |
| **FFA2** |  | 7.280 |  | 11 | |  | 1.583×10^-5^ |  | 2.102 |  | 0.517 |
| **People1** |  | 5.005 |  | 11 | |  | 3.992×10^-4^ |  | 1.445 |  | 0.413 |
| **People2** |  | 3.189 |  | 11 | |  | 0.009 |  | 0.921 |  | 0.344 |
| **Places1** |  | -5.813 |  | 11 | |  | 1.171×10^-4^ |  | -1.678 |  | 0.448 |
| **Places2** |  | -4.975 |  | 11 | |  | 4.185×10^-4^ |  | -1.436 |  | 0.411 |
| V1 0.969 11 0.177 0.280 0.294 | | | | | | | | | | | |
| *Note.*  For the Student t-test, effect size is given by Cohen's *d*. | | | | | | | | | | | |
| *Note.*  For the Student t-test, the alternative hypothesis specifies that the mean is different from 0. | | | | | | | | | | | |
| **Bayesian One Sample T-Test** | | | | | | | | | | | |
|  | | | | | **BF₁₀** | | | | | **error %** | |
| **PPAp** | | |  | | 7.576 | | | |  | 1.993×10^-6^ |  |
| **PPAa** | | |  | | 818.236 | | | |  | 7.623×10^-6^ |  |
| **FFA1** | | |  | | 260.685 | | | |  | 5.162×10^-6^ |  |
| **FFA2** | | |  | | 1441.540 | | | |  | 3.790×10^-6^ |  |
| **People1** | | |  | | 87.518 | | | |  | 1.838×10^-6^ |  |
| **People2** | | |  | | 6.674 | | | |  | 2.416×10^-6^ |  |
| **Places1** | | |  | | 251.548 | | | |  | 9.277×10^-7^ |  |
| **Places2** | | |  | | 84.069 | | | |  | 2.209×10^-6^ |  |
| V1 0.463 0.013 | | | | | | | | | | | |
| *Note.*  For all tests, the alternative hypothesis specifies that the population mean differs from 0. | | | | | | | | | | | |

**S5. MRI univariate analyses including hemisphere**

We entered the average beta values within each ROI for each subject into a linear mixed model with factors Hemisphere (left and right), Region (People1, People2, Places1, Places2), and Category (people and places), keeping the medial parietal and ventral temporal regions separate. Across both sets of ROIs, activation was largest for the preferred category. In the medial parietal regions, there was a significant main effect of Hemisphere (F(1,165) = 5.66, p = 0.018), driven by higher values in the left hemisphere (see **Figure S2**). There was also a significant main effect of ROI (F(3,165) = 17.36, p = 7.59e-10), a significant main effect of category (F(1,165) = 4.59, p = 0.034), and a significant interaction between Category and ROI (F(3,165) = 45.61, p = 2.2e-16).

In the ventral temporal regions (FFA1, FFA2, pPPA, aPPA), the main effect of Hemisphere was not significant (F(1,149.55) = 3.47, p = 0.064). There was a significant main effect of ROI (F(3,149.51) = 38.46, p = 2.2e-16), a significant main effect of Category (F(1,149.14) = 15.71 p = 0.001e-1), and a significant interaction between Category and ROI (F(3,149.14) = 45.60, p = 2.2e-16).

Next, we ran the same analysis using the univariate GLM beta values for people and place stimuli in V1. Here there were no significant effects of Category (F(1) = 1.15, p = 0.307), and no significant effects of Hemisphere (F(1) = 0.95, p = 0.444).

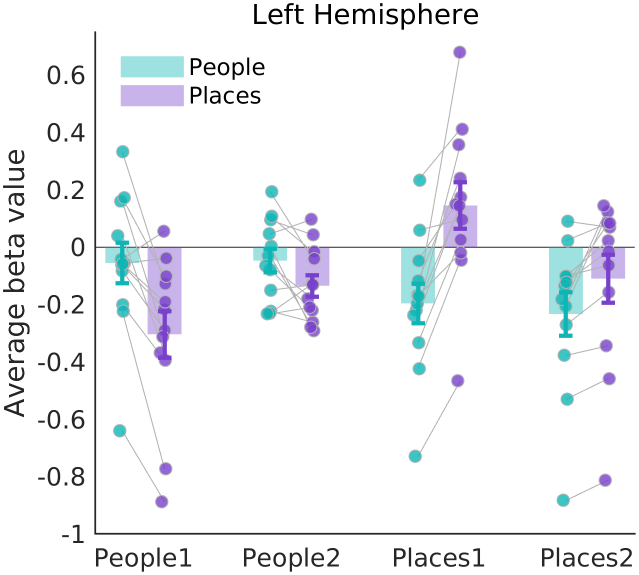

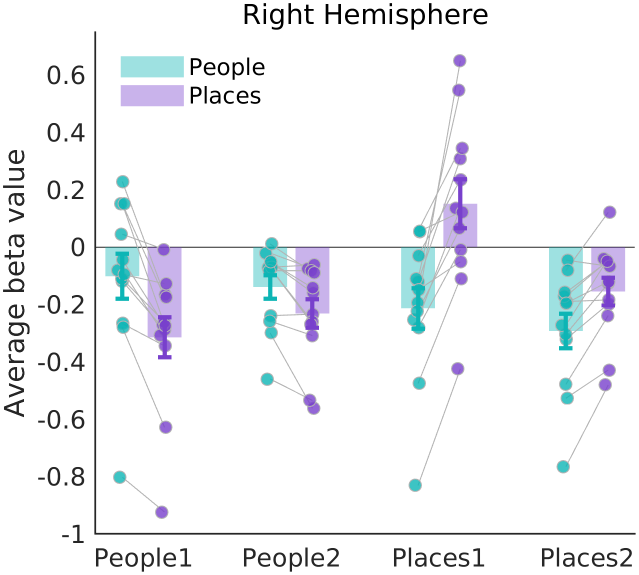

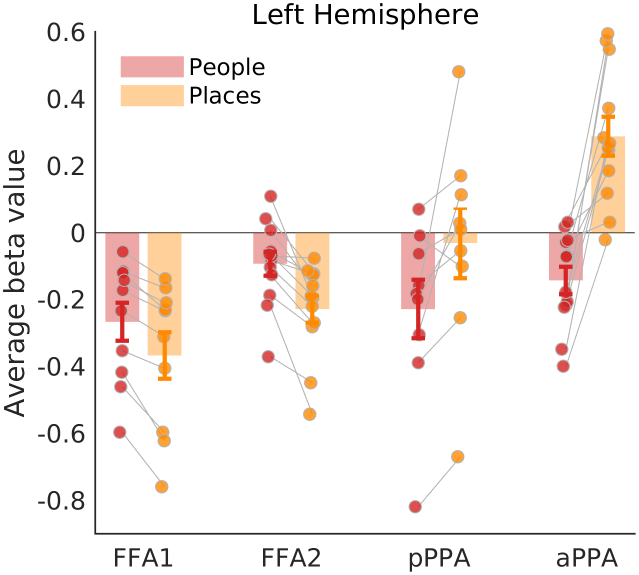

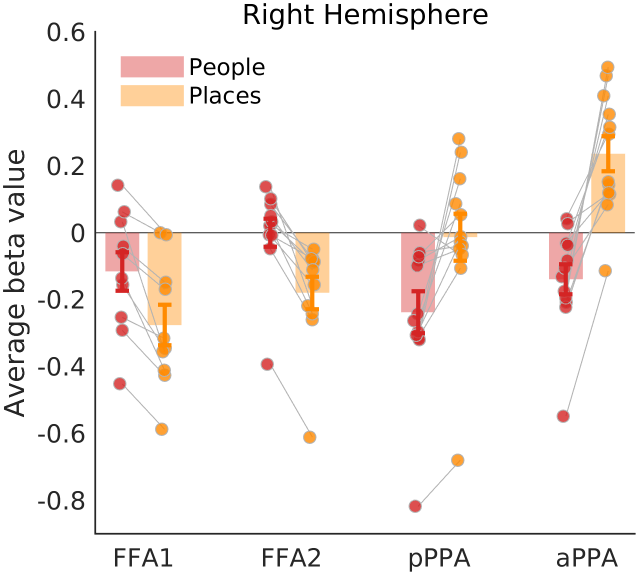

**Figure S2.** Average fMRI beta values separately for people and place imagery in medial parietal (top) and ventral temporal (bottom), separately for the left and right hemispheres. Each data point represents a single subject and error bars represent the standard error. FFA1 = posterior fusiform face area, FFA2 = anterior, pPPA = posterior parahippocampal place area, aPPA = anterior.

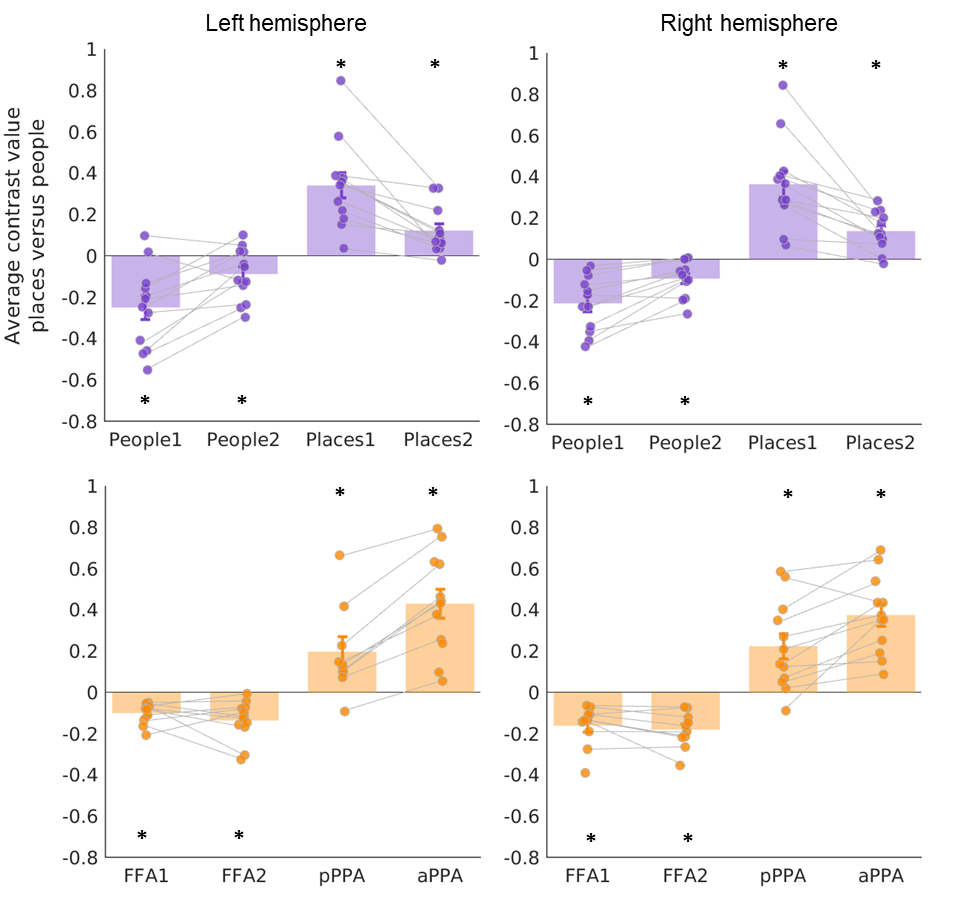
**Figure S3.** Average fMRI contrast values for places versus people imagery in the medial parietal (top) and ventral temporal cortex (bottom), separately for the left and right hemispheres. All contrast values were significantly different from zero, with values above zero in place selective regions and below zero in face selective regions (*p-values < .05). Each data point represents a single subject and error bars represent the standard error. FFA1 = posterior fusiform face area, FFA2 = anterior, pPPA = posterior parahippocampal place area, aPPA = anterior.

**S6. MRI region of interest univariate analysis: hippocampus and amygdala**

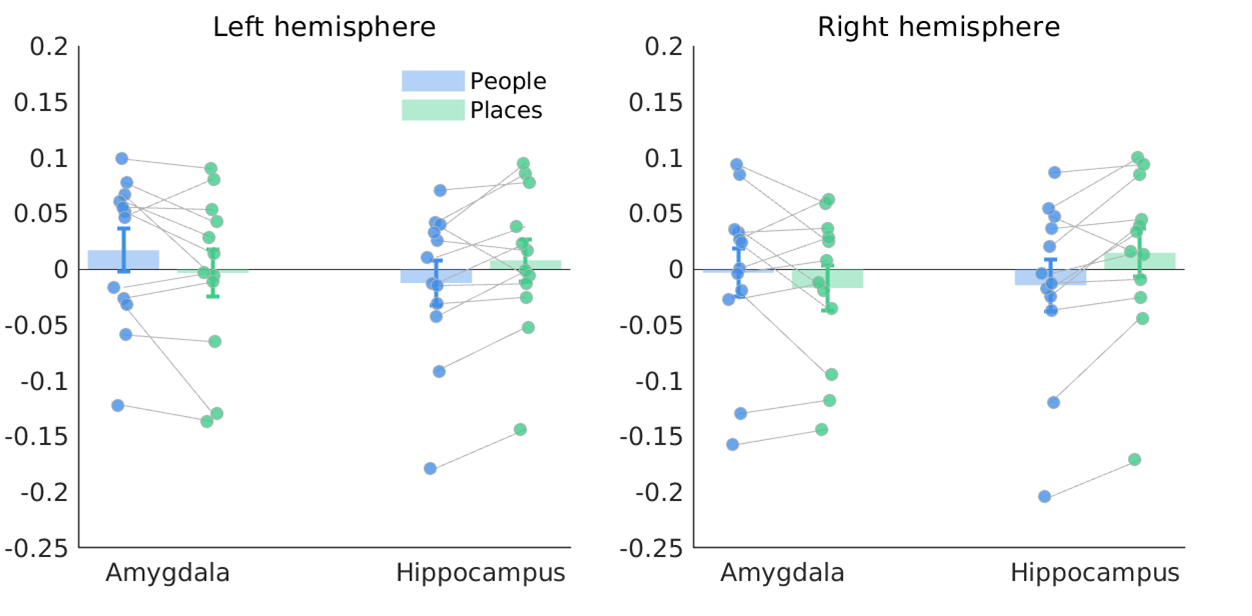
To replicate results from previous work (Silson et al., 2019), we also extracted data from the hippocampus and amygdala. We created masks using the Freesurfer Destrieux Atlas parcellations (aparc.a2009s) for each subject, which were aligned to the experiment structural (3dAllineate) and resampled to the same space as the pre-processed functional data (3dresample). We entered the average beta values for each subject into a repeated measures ANOVA with factors Region (hippocampus and amygdala), Hemisphere (left and right hemisphere), and Category (people and places). There was a significant interaction between Region and Category (F(11) = 12.049, p = 0.005). In one-tailed post-hoc t-tests, the contrast of places versus people was significantly greater than zero in the bilateral hippocampus (lh, t(11) = 2.399, p = .035; rh, t(11) = 3.128, p = .010). The contrast was significantly less than zero in the left amygdala only (lh, t(11) = -1.894, p = .042; rh, t(11) = -1.184, p = 0.131). The interaction was therefore driven by increased activation for places compared to people in the hippocampus, compared to the opposite effect in the left hemisphere amygdala. See **Figure S4** for the averaged beta values for each recall category.

**Figure S4:** Average fMRI beta values for places and people recall in the hippocampus and amygdala. The left and right hemispheres are displayed separately. Each data point represents a single subject and error bars represent the standard error.

**S7. MRI region of interest multivariate decoding: MPC and VTC**

**Category decoding:** results from one-sample Student and Bayesian t-tests comparing category decoding (person or place) in ventral temporal and medial parietal ROIs.

| **One Sample T-Test – decoding category** | | | | | | | | | | |
| --- | --- | --- | --- | --- | --- | --- | --- | --- | --- | --- |
|  | | **t** | | **df** | | **p** | | **Cohen's d** | | **SE Cohen's d** |
| **FFA1** |  | 3.898 |  | 10 |  | 0.003 |  | 1.175 |  | 0.392 |
| **FFA2** |  | 4.778 |  | 11 |  | 5.736×10^-4^ |  | 1.379 |  | 0.403 |
| **PPAp** |  | 5.317 |  | 11 |  | 2.461×10^-4^ |  | 1.535 |  | 0.426 |
| **PPAa** |  | 6.295 |  | 11 |  | 5.875×10^-5^ |  | 1.817 |  | 0.470 |
| **People1** |  | 6.675 |  | 11 |  | 3.493×10^-5^ |  | 1.927 |  | 0.488 |
| **People2** |  | 2.970 |  | 11 |  | 0.013 |  | 0.857 |  | 0.338 |
| **Places1** |  | 8.867 |  | 11 |  | 2.425×10^-6^ |  | 2.560 |  | 0.597 |
| **Places2** |  | 7.475 |  | 11 |  | 1.239×10^-5^ |  | 2.158 |  | 0.527 |
| *Note.*  For the Student t-test, effect size is given by Cohen's *d*. | | | | | | | | | | |
| *Note.*  For the Student t-test, the alternative hypothesis specifies that the mean is different from 0.5. | | | | | | | | | | |

| **Bayesian One Sample T-Test – decoding category** | | | | |
| --- | --- | --- | --- | --- |
|  | | **BF₁₀** | | **error %** |
| **FFA1** |  | 16.320 |  | 6.716×10^-6^ |
| **FFA2** |  | 64.231 |  | 4.265×10^-6^ |
| **PPAp** |  | 132.542 |  | 2.818×10^-4^ |
| **PPAa** |  | 457.807 |  | 3.175×10^-6^ |
| **People1** |  | 720.443 |  | 6.944×10^-6^ |
| **People2** |  | 4.860 |  | 4.080×10^-6^ |
| **Places1** |  | 7551.449 |  | 1.169×10^-8^ |
| **Places2** |  | 1788.171 |  | 3.784×10^-7^ |
| *Note.*  For all tests, the alternative hypothesis specifies that the population mean differs from 0.5. | | | | |

**Stimulus decoding:** results from one-sample Student and Bayesian t-tests comparing decoding of individual people and place stimuli in ventral temporal and medial parietal ROIs as plotted in **Figure 5** (FDR thresholds: MPC p < .022, VTC p < .019).

| **One Sample T-Test – decoding stimuli** | | | | | | | | | | | | |
| --- | --- | --- | --- | --- | --- | --- | --- | --- | --- | --- | --- | --- |
|  | | **t** | | **df** | | | **p** | | | **Cohen's d** | | **SE Cohen's d** |
| People1_people |  | 2.317 |  | 11 |  | | 0.041 | |  | 0.669 |  | 0.319 |
| **People1_places** |  | 3.245 |  | 11 |  | | 0.008 | |  | 0.937 |  | 0.346 |
| People2_people |  | 0.399 |  | 11 |  | | 0.697 | |  | 0.115 |  | 0.290 |
| **People2_places** |  | 2.853 |  | 11 |  | | 0.016 | |  | 0.824 |  | 0.334 |
| Places1_people |  | 1.830 |  | 11 |  | | 0.095 | |  | 0.528 |  | 0.308 |
| **Places1_places** |  | 6.718 |  | 11 |  | | 3.294×10^-5^ | |  | 1.939 |  | 0.490 |
| **Places2_people** |  | 2.666 |  | 11 |  | | 0.022 | |  | 0.770 |  | 0.329 |
| Places2_places |  | 1.892 |  | 11 |  | | 0.085 | |  | 0.546 |  | 0.309 |
| FFA1_people |  | 2.418 |  | 10 |  | | 0.036 | |  | 0.729 |  | 0.339 |
| FFA1_places |  | 1.432 |  | 10 |  | | 0.183 | |  | 0.432 |  | 0.315 |
| **FFA2_people** |  | 2.757 |  | 11 |  | | 0.019 | |  | 0.796 |  | 0.331 |
| FFA2_places |  | 2.516 |  | 11 |  | | 0.029 | |  | 0.726 |  | 0.325 |
| PPAp_people |  | 0.170 |  | 11 |  | | 0.868 | |  | 0.049 |  | 0.289 |
| **PPAp_places** |  | 5.046 |  | 11 |  | | 3.745×10^-4^ | |  | 1.457 |  | 0.414 |
| PPAa_people |  | 0.905 |  | 11 |  | | 0.385 | |  | 0.261 |  | 0.294 |
| **PPAa_places** |  | 5.055 |  | 11 |  | | 3.694×10^-4^ | |  | 1.459 |  | 0.415 |
| *Note.*  For the Student t-test, effect size is given by Cohen's *d*. | | | | | | | | | | | | |
| *Note.*  For the Student t-test, the alternative hypothesis specifies that the mean is different from 0.5. | | | | | | | | | | | | |
| *Note.*  Student's t-test. | | | | | | | | | | | | |
| **Bayesian One Sample T-Test – decoding stimuli** | | | | | | | | | | | | |
|  | | | | | | | | **BF₁₀** | | | **error %** | |
| People1_people | | | | | |  | | 1.939 | |  | 1.799×10^-6^ | |
| **People1_places** | | | | | |  | | 7.241 | |  | 2.103×10^-6^ | |
| People2_people | | | | | |  | | 0.308 | |  | 0.008 | |
| **People2_places** | | | | | |  | | 4.111 | |  | 4.371×10^-6^ | |
| Places1_people | | | | | |  | | 1.030 | |  | 0.023 | |
| **Places1_places** | | | | | |  | | 758.434 | |  | 7.262×10^-6^ | |
| **Places2_people** | | | | | |  | | 3.150 | |  | 3.273×10^-6^ | |
| Places2_places | | | | | |  | | 1.112 | |  | 0.024 | |
| FFA1_people | | | | | |  | | 2.189 | |  | 8.398×10^-7^ | |
| FFA1_places | | | | | |  | | 0.668 | |  | 0.017 | |
| **FFA2_people** | | | | | |  | | 3.584 | |  | 4.051×10^-6^ | |
| FFA2_places | | | | | |  | | 2.551 | |  | 1.418×10^-6^ | |
| PPAp_people | | | | | |  | | 0.291 | |  | 0.007 | |
| **PPAp_places** | | | | | |  | | 92.451 | |  | 1.356×10^-6^ | |
| PPAa_people | | | | | |  | | 0.406 | |  | 0.011 | |
| **PPAa_places** | | | | | |  | | 93.535 | |  | 1.260×10^-6^ | |
| *Note.*  For all tests, the alternative hypothesis specifies that the population mean differs from 0.5. | | | | | | | | | | | | |
| **Stimuli decoding:** paired t-tests comparing stimuli decoding in the preferred versus non-preferred category. **Paired Samples T-Test – decoding stimuli** \| \| \| \| \| \| \| \| \| \| \|; **Measure 1** \| \| \| \| **Measure 2** \| \| **t** \| \| **df** \| \| **p** \|; People1_people \| \| - \| \| People1_places \| \| -1.338 \| \| 11 \| \| 0.896 \|; People2_people \| \| - \| \| People2_places \| \| -1.328 \| \| 11 \| \| 0.895 \|; FFA1_people \| \| - \| \| FFA1_places \| \| 0.993 \| \| 10 \| \| 0.172 \|; **FFA2_people** \| \| **-** \| \| **FFA2_places** \| \| 2.058 \| \| 11 \| \| 0.032 \|; **PPAa_places** \| \| **-** \| \| **PPAa_people** \| \| 3.872 \| \| 11 \| \| 0.001 \|; **PPAp_places** \| \| **-** \| \| **PPAp_people** \| \| 3.374 \| \| 11 \| \| 0.003 \|; **Places1_places** \| \| **-** \| \| **Places1_people** \| \| 3.561 \| \| 11 \| \| 0.002 \|; Places2_places \| \| - \| \| Places2_people \| \| -0.280 \| \| 11 \| \| 0.608 \|; *Note.*  For all tests, the alternative hypothesis specifies that Measure 1 is greater than Measure 2. For example, People1_people is greater than People1_places. \| \| \| \| \| \| \| \| \| \| \|; *Note.*  Student's t-test. \| \| \| \| \| \| \| \| \| \| \| | | | | | | | | | | | | |

**S8. MRI region of interest multivariate decoding across hemispheres**

We entered the average decoding within each ROI into two linear mixed models with factors Hemisphere, Region, and Category, separately for the medial parietal and ventral temporal regions. For the ventral temporal cortex, the main effect of Hemisphere was not significant (F(1,150.92) = 1.02, p = 0.31). There was a significant interaction between ROI and Category (F(3,148.96) = 4.05, p = 0.008). The main effect of ROI was not significant (F(1,148.96) = 1.41, p = 0.24), and the main effect of category was also not significant (F(1,148.96) = 1.85, p = 0.18).

For the medial parietal cortex, the main effect of Hemisphere was not significant (F(1,165) = 0.99, p = 0.32). The interaction between ROI and Category was not significant (F(1,165) = 1.05, p = .31). There was a significant main effect of ROI (F(1,165) = 4.09, p = 0.008), and a significant main effect of Category (F(1,165) = 9.45, p = 0.002).

**S9. Fusion differences across hemisphere**

As some meta-analyses have indicated lateralised effects in perceptual and memory regions (Spagna et al., 2021; Winlove et al., 2018), we compared the fusion across left and right hemisphere ROIs. For each subject and ROI, we calculated the left minus right hemisphere fusion correlation timeseries. Across the group, we found Bayesian evidence predominantly in favour of the null hypothesis, indicating no difference in the fusion timeseries across hemispheres (see **Figure S5**).

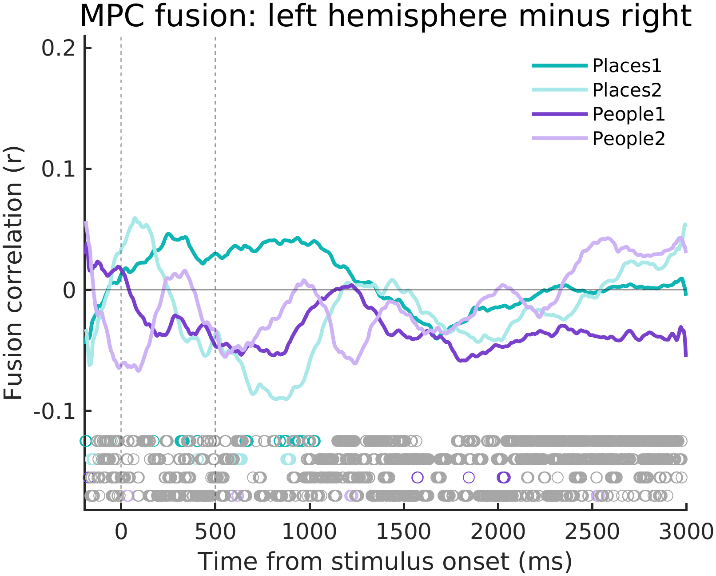

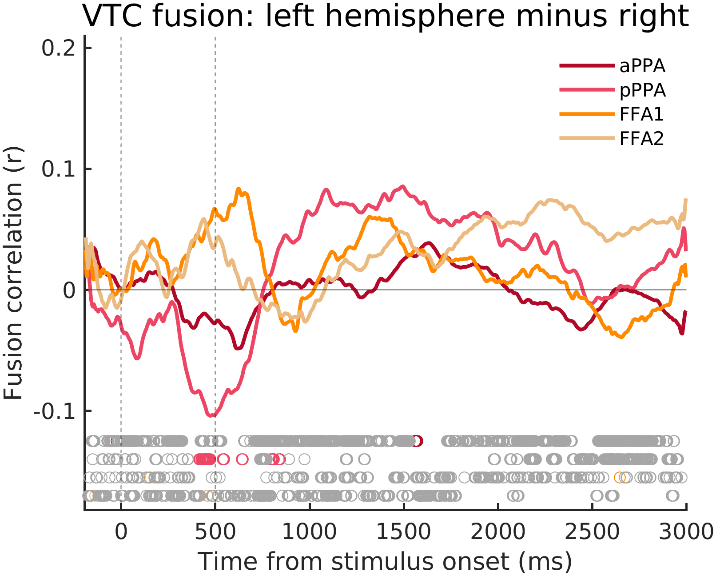

**Figure S5:** The difference in EEG-fMRI fusion timeseries across hemispheres (left minus right hemisphere). Medial parietal regions are plotted on the left-hand figure, and ventral temporal on the right-hand figure. Positive values indicate there was higher a fusion correlation in the left hemisphere, while negative values indicate that there was a higher fusion correlation in the right hemisphere. Timepoints with Bayes Factors that were greater than 3 are indicated by the coloured circles along the bottom bar, suggesting that the alternative hypothesis of a fusion correlation is at least three times more likely than the null hypothesis (BF_10_ >3). Grey circles indicate Bayes Evidence in favour of the null hypothesis (BF_10_ <1/3).

**S10. EEG and MRI fusion timeseries across hemispheres**

The following table contains the first time point with Bayesian evidence for the alternative hypothesis greater than three (BFalt > 3) for each ROI, as well as the time point with the highest fusion correlation value.

|  |  | Time from cue onset | Time from cue onset | |
| --- | --- | --- | --- | --- |
| **Hemisphere** | **ROI** | **Correlation onset (ms)** | | **Peak correlation (ms)** |
| Left | aPPA | 500 | | 727 |
|  | pPPA | 660 | | 730 |
|  | FFA1 | 949 | | 1379 |
|  | FFA2 | 2016 | | 2297 |
|  | Places1 | 535 | | 727 |
|  | Places2 | 586 | | 723 |
|  | People1 | 816 | | 930 |
|  | People2 | n/a | | n/a |
|  | V1 | 1426 | | 2094 |
| Right | aPPA | 500 | | 637 |
|  | pPPA | 543 | | 656 |
|  | FFA1 | 805 | | 871 |
|  | FFA2 | n/a | | n/a |
|  | Places1 | 543 | | 746 |
|  | Places2 | 500 | | 723 |
|  | People1 | 691 | | 832 |
|  | People2 | 609 | | 1738 |
|  | V1 | 1430 | | 1492 |

**S11. Peak locations from the whole brain univariate contrast of people versus place stimuli.** The peak locations of our ROIs were mapped from the MNI surface onto an MNI volume using the AFNI and SUMA viewers (LPI space). FFA = fusiform face area, VPMA = ventral place memory area, PPA = parahippocampal place area, LPMA = lateral place memory area, OPA = occipital place area, FEF = frontal eye fields, IPS = intraparietal sulcus.

| Region | Hemisphere | MNI (LPI) | Brodmann labels |
| --- | --- | --- | --- |
| FFA | LH | -37, -41, -22 | Left fusiform (37) |
|  | RH | 40, -45, -23 | Right fusiform (37) |
| VPMA/PPA | LH | -28, -38, -11 | Left parahippocampal (36) |
|  | RH | 31, -32, -15 | Right parahippocampal (36) |
| LPMA/OPA | LH | -37, -82, 28 | Left visual association cortex (19) |
|  | RH | 36, -80, 33 | Right angular gyrus (39) |
| FEF | LH | -21, 16, 52 | Left premotor and supp. motor (6) |
|  | RH | 29, 10, 45 | Right frontal eye fields (8) |
| IPS | LH | -10, -77, 54 | Left superior parietal lobule/precuneus (7) |
|  | RH | 5, -68, 53 | Right superior parietal lobule/precuneus (7) |
| Inferior temporal | LH | -51, -47, -12 | Left fusiform (37) |
|  | RH | 52, -53, -11 | Left fusiform (37) |
| Post central | LH | -42, -30, 37 | Left supramarginal gyrus (40) |
|  | RH | 35, -35, 38 | Right supramarginal gyrus (40) |
| Superior temporal sulcus | RH | 39, -55, 24 | Right angular gyrus (39) |
| Places 1 | LH | -11, -55, 11 | Left ventral posterior cingulate (23) |
|  | RH | 16, -54, 14 | Right ventral posterior cingulate (23) |
| Places 2 | LH | -6, -38, 44 | Left dorsal posterior cingulate (31) |
|  | RH | 9, -43, 50 | Right dorsal posterior cingulate (31) |
| People 1 | LH | -13, -43, 32 | Left ventral posterior cingulate (23) |
|  | RH | 7, -53, 28 | Right ventral posterior cingulate (23) |
| People 2 | LH | -4, -14, 36 | Left ventral posterior cingulate (23) |
|  | RH | 6, -18, 36 | Right ventral posterior cingulate (23) |
